# Conditional deletion of CEACAM1 causes hepatic stellate cell activation

**DOI:** 10.1101/2024.04.02.586238

**Authors:** Harrison T. Muturi, Hilda E. Ghadieh, Suman Asalla, Sumona G. Lester, Stefaan Verhulst, Hannah L. Stankus, Sobia Zaidi, Raziyeh Abdolahipour, Getachew D. Belew, Leo A van Grunsven, Scott L. Friedman, Robert F. Schwabe, Terry D. Hinds, Sonia M. Najjar

## Abstract

**Objectives:** Hepatic CEACAM1 expression declines with advanced hepatic fibrosis stage in patients with MASH. Global and hepatocyte-specific deletions of *Ceacam1* impair insulin clearance to cause hepatic insulin resistance and steatosis. They also cause hepatic inflammation and fibrosis, a condition characterized by excessive collagen production from activated hepatic stellate cells (HSCs). Given the positive effect of PPARγ on CEACAM1 transcriptoin and on HSCs quiescence, the current studies investigated whether CEACAM1 loss from HSCs causes their activation.

**Methods:** We examined whether lentiviral shRNA-mediated CEACAM1 donwregulation (KD-LX2) activates cultured human LX2 stellate cells. We also generated *LratCre+Cc1^fl/fl^* mutants with conditional *Ceacam1* deletion in HSCs and characterized their MASH phenotype. Media transfer experiments were employed to examine whether media from mutant human and murine HSCs activate their wild-type counterparts.

**Results:** *LratCre+Cc1^fl/fl^* mutants displayed hepatic inflammation and fibrosis but without insulin resistance or hepatic steatosis. Their HSCs, like KD-LX2 cells, underwent myofibroblastic transformation and their media activated wild-type HDCs. This was inhibited by nicotinic acid treatment which stemmed the release of IL-6 and fatty acids, both of which activate the epidermal growth factor receptor (EGFR) tyrosine kinase. Gefitinib inhibition of EGFR and its downstream NF-κB/IL-6/STAT3 inflammatory and MAPK-proliferation pathways also blunted HSCs activation in the absence of CEACAM1.

**Conclusions:** Loss of CEACAM1 in HSCs provoked their myofibroblastic transformation in the absence of insulin resistance and hepatic steatosis. This response is mediated by autocrine HSCs activation of the EGFR pathway that amplifies inflammation and proliferation.

## 1. Introduction

Metabolic-Associated Steatotic Liver Disease (MASLD), formerly termed non-alcoholic fatty liver disease, currently represents the most common cause of chronic liver disease worldwide [1]. MASLD spans a broad spectrum of metabolic disease with hepatic fibrosis defining its most aggressive form, Metabolic-Associated Steatohepatitis (MASH), together with inflammation, hepatocyte damage, and apoptosis. Hepatic fibrosis is on the rise and currently constitutes a leading etiology in patients with MASH, partly because of limited targeted therapy [2; 3]. This necessitates the need for further studies exploring its molecular and cellular basis.

Histologically, hepatic fibrosis in patients with MASH is characterized by early lesions of perisinusoidal collagen deposition, followed by portal and eventually, bridging fibrosis [4]. It implicates the activation of hepatic stellate cells (HSCs) located in the Space of Disse between liver sinusoidal endothelial cells (LSECs) and hepatocytes. HSCs represent approximately 10% of resident liver cells. In healthy liver, PPARγ activation maintains HSCs quiescent and containing large lipid droplets filled with vitamin A as retinyl esters (RE), triacylglycerols (TG) and cholesteryl esters (CE) [5]. Following transdifferentiation into proliferative, contractile, inflammatory myofibroblasts with enhanced extracellular matrix (ECM) production, HSCs lose their retinoid content [6]. This is associated with reduced PPARγ and reciprocal elevation in the level of PPARβ/δ [7; 8], which could be activated by all trans-retinoic acid [9] and PUFA [10] to increase HSCs proliferation via inducing the p38 and JNK MAPK pathways [8]. Further studies are needed to fully identify the factors that cause HSCs activation [11].

Virtually every liver cell contributes to HSCs activation, and they all express the Carcinoembryonic Antigen-related Cell Adhesion Molecule 1 (CEACAM1), with a dominant expression in hepatocytes where it promotes insulin clearance. Depletion of *Ceacam1* gene globally [12] or exclusively in hepatocytes [13], causes chronic hyperinsulinemia, emanating chiefly from reduced insulin clearance, followed by hepatic insulin resistance, steatohepatitis and visceral obesity. It also provokes HSCs activation and a characteristic MASH-like fibrosis [14; 15]. Fed a high-fat diet, mice lacking CEACAM1 in hepatocytes develop advanced hepatocellular injury accompanied by chicken-wire fibrosis and apoptosis [14; 16]. Reciprocally, liver-specific rescuing of CEACAM1 reverses metabolic dysregulation and hepatic fibrosis in global *Cc1^−/−^* null mice [14]. In contrast, CEACAM1 loss in endothelial cells promotes hepatic fibrosis, driven by increased production of endothelin1, without insulin resistance or hepatic steatosis [17]. Consistent with these data in genetically-modified mice, patients with MASH exhibit a progressive loss of CEACAM1 in liver [15] and particularly in LSECs [17] as the disease advances.

CEACAM1 is also expressed in pericytes [18], including HSCs [19]. Herein we investigated whether its loss of CEACAM1 in HSCs induces their activation and sought to uncover underlying mechanisms.

## 2. Materials and methods

### 2.1. Generation and metabolic phenotyping of LratCre+Cc1^fl/fl^ mice

As detailed in Supplemental data, *Cc1loxp/loxp* mice were crossed with *LratCre* transgenic mice expressing a Cre recombinase driven by mouse lecithin-retinol acyltransferase (Lrat) promoter [20]. Stellate cell-specific deletion of *Ceacam1* in C57BL/6Jxhomozygotes (*LratCre+Cc1^fl/fl^*) was confirmed by PCR reaction using gene-specific primers (Fig. S1). As littermate controls, this study used homozygotes of wild-type *Ceacam1* allele with (*LratCre+Cc1^+/+^*) or without *Cre* (*LratCre–Cc1^+/+^*), and homozygotes of *Ceacam1*-floxed allele without *Cre* (*LratCre–Cc1^fl/fl^*) to rule out potential confounding effects of floxing and introducing *Cre* recombinase.

Per institutionally approved protocols, animals were housed in a 12-h dark-light cycle and fed standard chow *ad libitum*. Male mice were kept in cages with Alpha-dri bedding before undergoing metabolic phenotyping [intraperitoneal (IP) glucose and insulin tolerance tests-GTT and ITT, respectively]. Following recovery, mice were fasted for 18hrs, anesthetized with an IP injection of pentobarbital (1.1mg/kg BW), and their retro-orbital venous blood was drawn and tissues extracted for biochemical evaluation (Supplemental data).

### 2.2. Liver histology and immunohistochemical analysis

As detailed in Supplemental data, fixed liver sections were stained with hematoxylin-eosin (H&E) or with 0.1% Sirius Red stain to evaluate hepatic fibrosis. Images were taken using Nikon Eclipse 90i Microscope and 10 randomly selected high power fields (20X) per sample were imaged with ImageJ (v1.53t) to quantify Sirius Red stain as %area [17].

For immunohistochemical (IHC) analysis, liver sections underwent antigen-retrieval, blocking with rabbit or mouse serum, stained overnight at 4°C with specific antibodies, blotted with species-specific biotinylated secondary antibodies before being hematoxylin-counterstained [17]. Images were taken using Nikon Eclipse 90i Microscope and evaluated blindly to count positively stained cells in 5 fields/mouse at 40X magnification.

### 2.3. Isolating HSCs from human subjects

Human HSCs were isolated, as described [21]. Briefly, dissociated parenchymal cells were suspended in 2mM EDTA buffer containing 5%FBS (Gibco) for 30min in the presence of anti-CD32 (Abcam, Cambridge, MA) and anti-CD45 (BD Biosciences, San Jose, CA). Quiescent HSCs (qHSCs) were sorted as CD32-CD45-UV+ cells using FASCAria (BD Biosciences). Activated HSCs (aHSCs) were obtained by plating qHSCs in DMEM (Gibco) supplemented with 20% FBS, 20ng/mL epidermal growth factor (EGF) and 10ng/mL fibroblast growth factor 2 (Peprotech, London, UK), 100μM oleic acid, 100μM palmitic acid, and 5μM retinol (Sigma-Aldrich). RNA was isolated using RNeasy Micro Kit (Qiagen GmbH, Hilden, Germany). RNA samples were amplified using the Ovation Pico WTA system V2 (Tecan Genomics, San Carlos, CA).

### 2.4. Media transfer experiments in primary murine hepatic stellate cells

Primary HSCs were isolated from ≥8-month-old control *LratCre–Cc1^fl/fl^* (recipient cells) and mutant mice *LratCre+Cc1^fl/fl^* (donor cells) [22]. Cells were cultured in 12-well-plates for 5 days. *LratCre+Cc1^fl/fl^* donor cells were washed twice and incubated in phenol red-free DMEM before treating with 500µM nicotinic acid (NA) (Sigma-Aldrich) or buffer alone for 24hrs. Media were collected, centrifuged at 380g for 3min to remove cell debris and the “conditioned media” were transferred to the twice-washed *LratCre– Cc1^fl/fl^* recipient HSCs. 24hrs later, cells were lysed for mRNA analysis (see below). In some experiments, 10µM Gefitinib (Sigma-Aldrich), an EGFR tyrosine kinase inhibitor [23], or dimethyl sulfoxide (DMSO-vehicle) were added to recipient cells for additional 24hrs before cell lysis. Media levels of free glycerol (Glycerol Assay Kit MAK117-1KT, Sigma-Aldrich), interleukin-6 (ELISA Kit, ab222503, Abcam) and TNFα (ELISA Kit, ab100747, Abcam) were determined per manufacturer instructions [17].

### 2.5. Experiments with LX2 cells with stable downregulation of human CEACAM1 expression

The immortalized human hepatic stellate LX2 cell line was infected with a human CEACAM1 shRNA lentiviral construct to establish a KD line with stable knockdown of hCEACAM1 and scramble control (Scr), as detailed in Supplemental data. KD-LX2 and Scr-LX2 cells were treated with DMSO, 5μM retinoic acid and/or 1μM Rosiglitazone (Sigma-Aldrich) for 24hrs before cell lysis and qRT-PCR analysis [16].

For lipid analysis, KD-LX2 and Scr-LX2 cells were seeded in 6-well-plates (4x10^4^ cells/well) for 48hrs before being stained with Nile Red (Sigma-Aldrich) and evaluated with densitometry by ImageJ software to measure lipid content [24]. Media was collected to determine free glycerol levels using Glycerol Assay Kit (BioVision, Milpitas, CA) [24].

Media transfer from KD-LX2 to Scr-LX2 controls was performed as above and levels of free glycerol (MAK117-1KT, Sigma-Aldrich), interleukin-6 (ELISA-ab 178018, Abcam) and TNFα (ELISA-ab181421, Abcam) were determined per manufacturer instructions.

Cell growth was determined by MTT assay (Sigma-Aldrich) and absorbance read at 570nm in 96-well plates. Cell growth was calculated as percent of growth in the presence of effector minus basal growth divided by maximum growth in complete medium.

### 2.6. Immunoprecipitation and Western blot analysis

As previously described [17], cells were Triton-lysed and subjected to SDS-PAGE followed by Western blot analysis using antibodies as listed in Supplemental data. Proteins were detected by chemiluminescence, scanned and their density normalized against tubulin (Cell Signaling) or the total amount of proteins of the signaling molecule applied on parallel gels.

For immunoprecipitation, 100µg of protein lysates were precleared with 20µl mixture of protein G and A sepharose beads (Invitrogen, Carlsbad, CA) at 4^0^C for 2hrs. Proteins were immunoprecipitated from the precleared lysates by incubation with 2µg of their specific antibodies overnight at 4^0^C, centrifuged and analyzed by SDS-PAGE and Western blot analysis.

### 2.7. Quantitative real-time-PCR (qRT-PCR)

Total RNA was isolated with PerfectPure RNA Tissue Kit (Fisher Scientific, Waltham, MA). cDNA was synthesized by iScript cDNA Synthesis Kit (Bio-Rad), using 1μg of total RNA and oligodT primers (Table S1). cDNA was evaluated with qRT-PCR (StepOne Plus, Applied Biosystems, Foster City, CA), and mRNA was normalized to *GAPDH*, unless otherwise mentioned.

### 2.8. Statistical analysis

Data were analyzed using one-way ANOVA analysis with Bonferroni correction or two-tailed Student-t-test using GraphPad Prism 6 software. Data were presented as means±SEM. *P*<0.05 was considered statistically significant.

## 3. Results

### 3.1. CEACAM1 expression is induced by rosiglitazone and retinoic acid

Rosiglitazone (Ro) elevated *CEACAM1* mRNA levels by ∼2-to-3–fold in human LX2 HSCs (Fig. 1A). This likely stems from the transcriptional activation of *Ceacam1* promoter by the binding of liganded PPARγ to the functional and well-conserved PPAR response element/retinoic acid receptor recognition site (PPRE/RXRα) between nts–557 and –543 in *Ceacam1* promoter [25]. A similar effect was exerted by retinoic acid (RA) either alone or combined with rosiglitazone (Fig. 1A). This resulted from the transcriptional activation by RA, as tested by a luciferase reporter assay in human HepG2 cells transfected with mouse *Ceacam1* promoter [25] (Fig. 1B). As shown, Ro and RA induced the luciferase activity of PPREx3TK-Luc and PGL3-RARE-Luc (positive controls for Ro and RA, respectively) and of the mouse *Ceacam1* promoter (–1100pLuc), individually or combined, without affecting that of the empty vector (PGL4.10) [Fig. 1B, \**P*<0.05 versus vehicle-treated (–)]. Mutating the PPRE/RXRα sequence (–Δ2), PPRE (–ΔPPRE) or RXRα (–ΔRXRα) abolished the positive effect of Ro and RA on *Ceacam1* promoter activity.

**Fig. 1.**
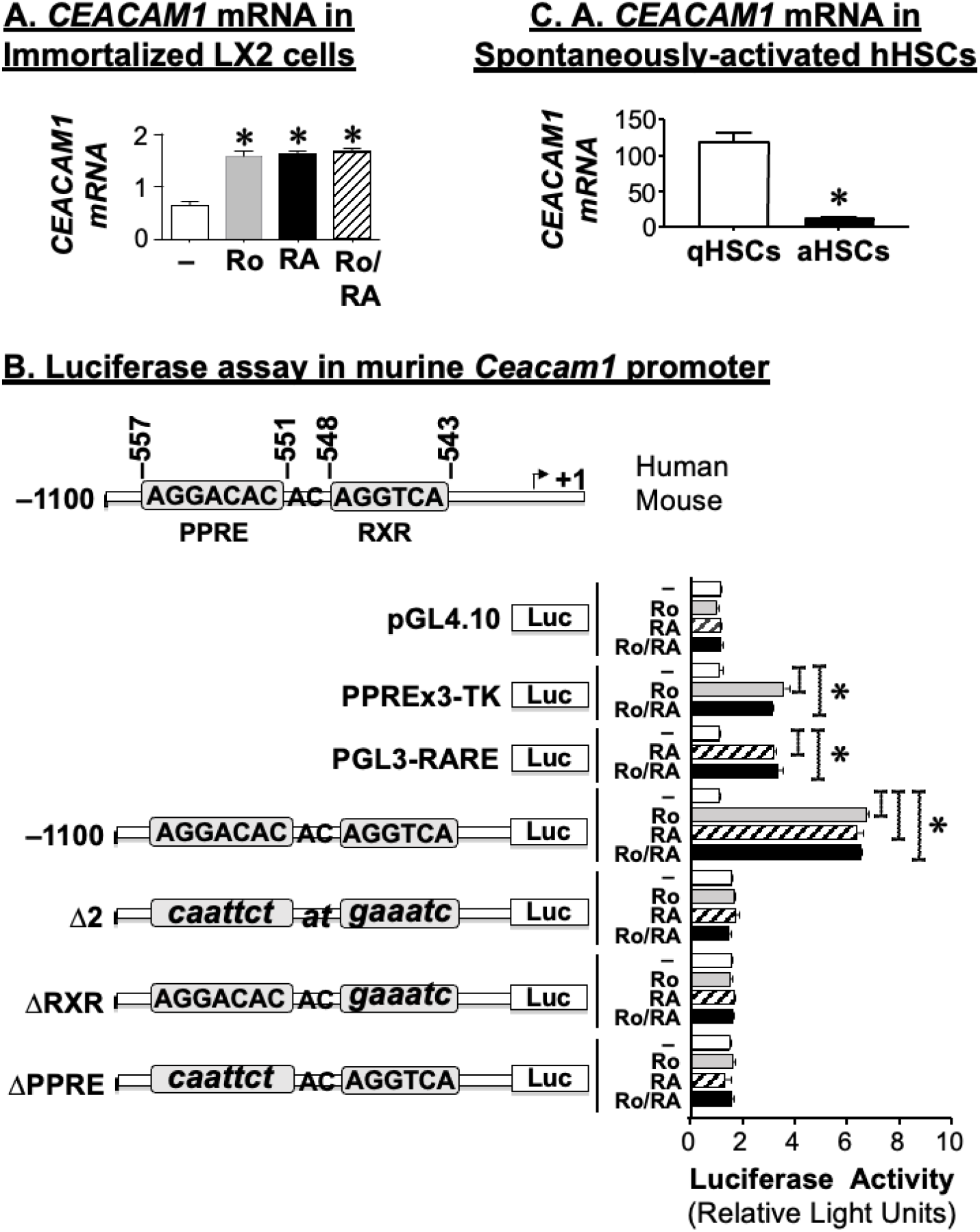
Regulation of CEACAM1 transcription. (A) immortalized human hepatic stellate cells LX2 cells were treated with DMSO (–) (white bars), 1 μM Rosiglitazone (Ro) (grey bars), 5 μM Retinoic Acid (RA) (black bars), and Ro plus RA (hatched bars) for 24 hrs before being subjected to qRT-PCR analysis of *CEACAM1* (*CC1*) mRNA levels. Data are expressed as mean ± SEM; \**P*<0.05 vs vehicle (–). (B) to analyze the transcriptional regulation of Ceacam1 promoter activity in HepG2, wild-type (nts −1100) mouse *Ceacam1* promoter and block mutants (small letters) of the PPRE/RXRα site (nts –557 to –543) (–Δ2); RXRα (–548 and –543) (–ΔRXRα); or PPRE (nts –557 and -551) (–ΔPPRE) were subcloned into pGL4.10 promoterless plasmid. Luciferase activity was measured in triplicate in response to DMSO (–, white), rosiglitazone (Ro, grey), retinoic acid (RA, checkered) or rosiglitazone plus retinoic acid (Ro/RA, black). PPREx3-TK-luc and PGL3-RARE-Luc were used as positive controls for PPRE and RXR, respectively. PGL4.10 empty vector was used as a negative control. Luciferase light units are expressed as mean ± SEM in relative light units. **\****P*<0.05 treatment versus vehicle. (C) *CEACAM1* mRNA was evaluated in spontaneously activated (aHSCs) (black bars) and quiescent (qHSCs) (white bars) primary human HSCs. Data are expressed as mean ± SEM; * *P*<0.05 vs qHSCs.

### 3.2. Loss of CEACAM1 activates human LX2 stellate cells

Activated primary human HSCs (aHSCs) exhibited lower (by >80%) CEACAM1 mRNA levels relative to quiescent cells (qHSCs) (Fig. 1C). To test whether CEACAM1 loss mediated HSC activation, we examined whether lentiviral shRNA-mediated repression of CEACAM1 by >90% (Fig. 2A.i and ii) could activate LX2-HSCs. Consistent with the loss of lipid content during HSC activation [26–28], knocking down CEACAM1 markedly reduced Nile red-stained fat-laden droplets relative to scrambled controls (Fig. 2B.i KD vs Scr in the graphical presentation of densitometry analysis). The lost cellular fat was recovered as free glycerol in the KD-LX2 culture media (Fig. 2B.ii). As predicted based on the known features of HSC activation, KD-LX2 cells exhibited reduced retinyl ester (RE) synthesis, as assessed by lower mRNA levels of enzymes catalyzing RE synthesis [lecithin-retinol acyltransferase (LRAT)] and lipolysis [lysosomal acid lipase-LAL (LIPA) [29]] (Fig. 2B.iii). Reciprocally, they manifested elevated PUFA-triacylglycerol (PUFA-TG) synthesis [27], as suggested by their higher ratio of the mRNA of PUFA-specific fatty acid-CoA synthase 4/non-specific ACSL1 (ACSL4)/ACSL1 [30], and their two-to-threefold higher mRNA levels of lipogenic genes such as SREBP-1c and fatty acid synthase (FASN), DGAT1 (the last enzyme in TG synthesis) and ATGL (TG lipase).

**Fig. 2.**
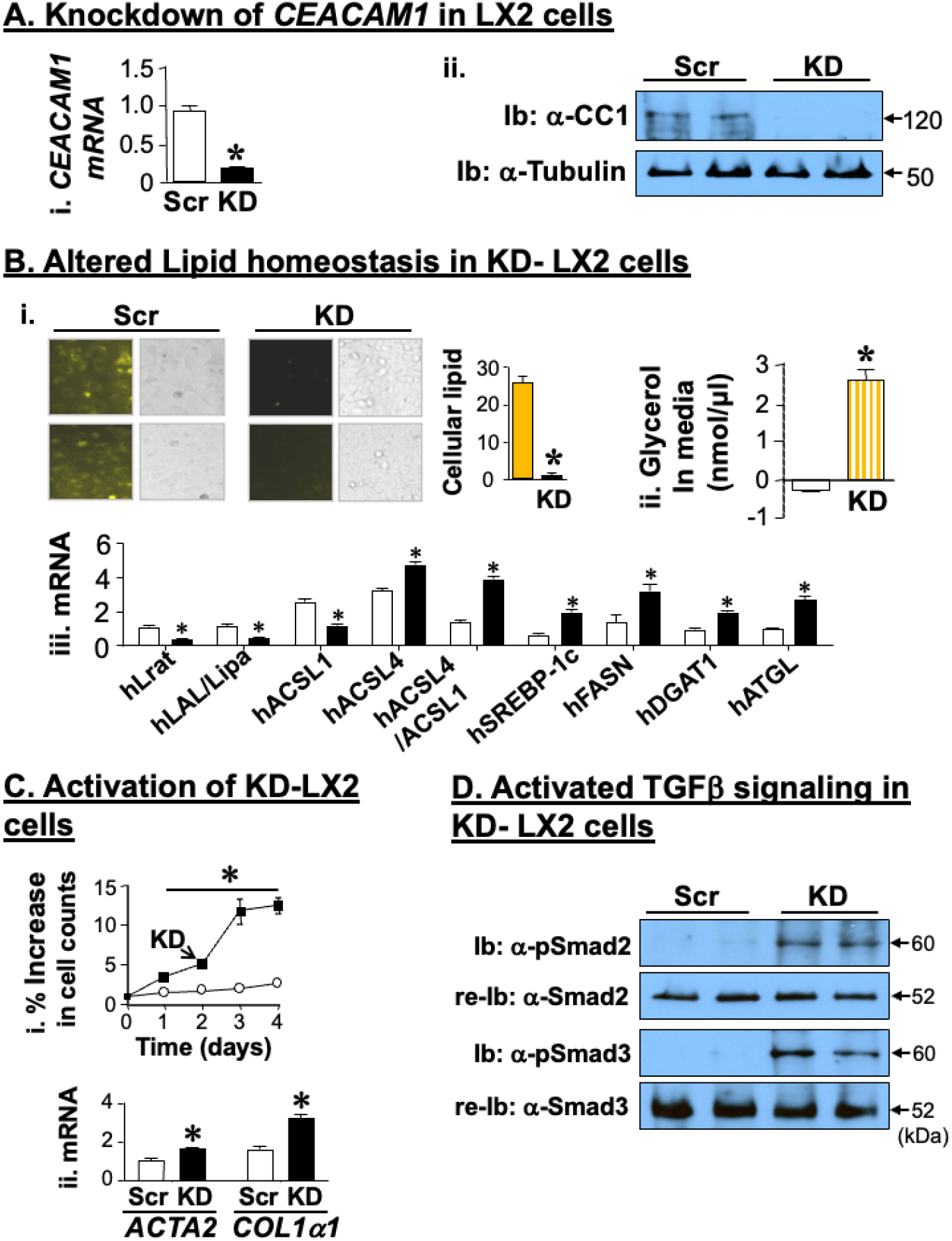
Activation of LX2 human hepatic stellate cells by CEACAM1 deletion. (A) LX2 cells were subjected to shRNA-mediated knockdown of *CEACAM1* (KD) and (i) analyzed by qRT-PCR in triplicate to assess the decrease in *CEACAM1* mRNA in KD-LX2 (Black bars) versus Scr-LX2 scrambled control cells (white bars). mRNA was normalized to *GAPDH* mRNA and data represented as mean ± SEM; * *P*<0.05 vs Scr-LX2; (ii) CEACAM1 protein levels were assessed by immunoblotting (Ib) the upper half of the membrane with α-CEACAM1 (α-CC1) antibody and the lower half with α-tubulin to normalize per loaded proteins. (B) to examine lipid metabolism, (i) cells were grown in at least 3 plates/stable line, stained with Nile Red to depict fat (yellow) droplets. Fat stains were evaluated by densitometry and presented graphically as mean ± SEM; * *P*<0.05 vs Scr-LX2; (ii) free glycerol level was assayed in the media of the stained cells as a measure of lipolysis. Experiments were done in triplicate. Data are expressed as mean ± SEM; * *P*<0.05 vs Scr-LX2; (iii) qRT-PCR mRNA analysis of genes implicated in lipid metabolism was performed in triplicate. Values are expressed as mean ± SEM. \**P*< 0.05 vs Scr-LX2. (C) to assess LX2 activation, (i) KD-LX2 and Scr-LX2 cells were subjected to MTT assay in triplicate. Data represent mean ± SEM; \**P*<0.05 vs Scr-LX2; (ii) qRT-PCR analysis was performed in triplicate to assess *ACTA2* and *COL1α1* mRNA levels as markers of fibrogenic activity. Data represent mean ± SEM; * *P*<0.05 vs Scr-LX2. (D) to examine TGFβ signaling, cell lysates were immunoblotted with α-phosphoSmad2 or α-phosphoSmad3 antibody (α-pSmad) followed by re-immunoblotting (re-Ib) with α-Smad 2 or α-Smad 3 antibodies, respectively, for normalization.

Activated HSCs undergo proliferation and resist apoptosis [26; 31]. Consistently, knocking down CEACAM1 markedly increased LX2 proliferation, as assessed by MTT assay (Fig. 2C.i). It also led to ∼2-fold higher mRNA levels of α-smooth muscle actin (*α-SMA or ACTA2*) and *COL1α1*, markers of mesenchymal cell activation (myofibroblastic transformation) [26] (Fig. 2C.ii). The latter could result from increased activation of TGFβ canonical signaling pathway, as assessed by Western blot analysis of phosphorylated Smad2/3 (Fig. 2D). This demonstrated that CEACAM1 loss activated KD-LX2 cells.

### 3.3. Delineating the mechanism underlying LX2 activation by CEACAM1 deletion

Following phosphorylation by epidermal growth factor (EGFR) and insulin (IR) receptors, CEACAM1 sequesters Shc and reduces its coupling to the receptors to suppress downstream Shc/MAPK-mediated cell growth and proliferation pathways [32; 33]. Consistently, insulin (100 nM) treatment for 5 min stimulated IRβ and MAPK phosphorylation in both groups of cells (Fig. S2B-C, + vs – insulin), and induced their proliferation, as assessed by MTT assay (Fig. S2D, + vs – insulin). In the absence of insulin, KD-LX2 cells manifested a higher basal phosphorylation of MAPK, but not IRβ (Fig. S2B-C), in parallel to higher cell growth relative to their Scr-LX2 controls (Fig. S2D). This suggests that an IR-independent pathway was implicated in KD-LX2 basal activation.

Because lipolysis-derived FAs activate EGFR [33], which contributes to HSC activation [34], we then examined whether the FAs released from KD-LX2 cells [and captured as free glycerol in their media (Fig. 3A.i, black vs white bar)], could activate EGFR pathways to induce HSC myofibroblastic transformation. As Fig. 3B.i shows, incubating Src-LX2 in the conditioned media of KD-LX2 cells (Scr/Cond) markedly reduced *CEACAM1* (*CC1*) mRNA levels (∼75%) (grey vs white bar; *P*<0.05). This likely resulted from increased *PPARβ/δ* expression (Fig. 3B.ii) and its activation by the released FAs. Consistently, blocking lipolysis by nicotinic acid (NA) normalized FAs content in KD-LX2 media (Fig. 3A.i, diagonally hatched vs white and vertically-hatched bars) and subsequently, restored *CEACAM1* mRNA levels in Scr/Cond (Figs. 3B.i, horizontally hatched vs vertically hatched bar).

**Fig. 3.**
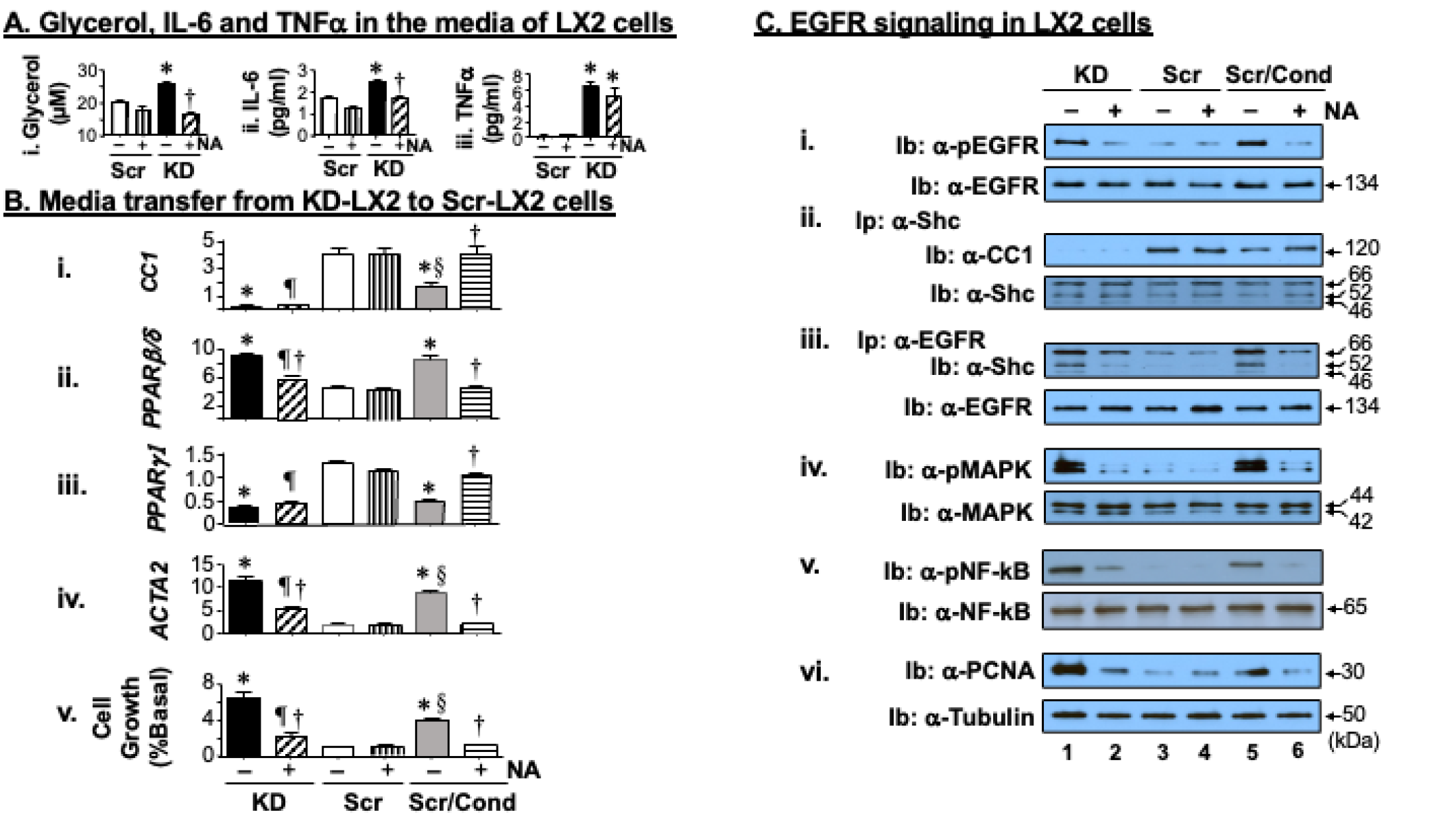
Scr-LX2 activation by conditioned media from KD-LX2 cells. Src-LX2 and KD-LX2 cells were incubated with nicotinic acid (NA) (+) or with vehicle (–) for 24 hrs before (A) media were collected and assayed for glycerol (captured lipolysis-derived fatty acids) (i), IL-6 (ii) and TNFα (iii) levels. Data represent mean ± SEM; \**P*<0.05, KD-LX2 vs Scr-LX2 cells/treatment type; ^†^p<0.05, NA-treated vs untreated/cell line. (B) media of KD-LX2 cells (conditioned media) were transferred to pre-washed Scr group (Scr/Cond) before cells were harvested for qRT-PCR analysis of the mRNA of *CC1*, *PPARβ/δ, PPARγ1 and ACTA2* (Fig. 3B.i-iv) and cell growth by MTT assay in triplicate and repeated twice (Fig. 2B.v). \**P*<0.05 untreated KD and Scr/Cond vs untreated Scr-LX2 cells; ^†^*P*<0.05 NA-treated vs untreated/cell line; ^§^*P*<0.05 untreated Scr/Cond vs untreated KD, and ^¶^*P*<0.05 NA-treated KD vs NA-treated Scr and NA-treated Scr/Cond cells. The latter indicates that although NA treatment decreased the mRNA of *PPARβ/δ* and *ACTA2* as well as cell growth, it did not completely restore their values to those in Scr-LX2 controls, as it did in Scr/Cond cells. This is likely due to persistent absence of CEACAM1 (and low *PPARγ1*) with sustained TNFα (iii) levels in these donor KD-LX2 cells. (C) Western blot analysis of EGFR signaling in cells described above: liver lysates were subjected to immunoblotting (Ib) with antibodies against (i) phospho-EGFR (α-pEGFR), (iv) phospho-MAPK (α-pMAPK), (v) phospho-NF-kB (α-pNF-kB), and (vi) α-PCNA and in parallel gels, with their specific antibodies for normalization. (ii) some lysates were subjected to immunoprecipitation (Ip) with Shc antibody followed by immunoblotting (Ib) with antibodies against CEACAM1 (α-CC1) and Shc (α-Shc). (iii) Lysates were subjected to immunoprecipitation (Ip) with α-pEGFR antibody followed by immunoblotting (Ib) with α-Shc and α-EGFR antibodies. Gels represent two separate experiments. The apparent molecular mass (kDa) is indicated at the right hand-side of each gel.

Western blot analysis revealed higher EGFR phosphorylation in KD and Src/Cond cells in the absence of NA (Fig. 3C.i, – lanes 1 and 5 vs lane 3), but not in its presence (Fig. 3C.i, + vs – lanes/cell group). This demonstrated that EGFR was activated in response to FA-containing conditioned media. Consistent with reduced CEACAM1 level, CEACAM1/Shc binding was lower in Scr/Cond than Scr cells, as demonstrated by its repressed detection in the Shc immunopellet (Fig. 3C.ii, – lane 5 in Scr/Cond vs – lane 3 in Scr). This led to a reciprocal recovery of Shc in the EGFR immunopellet of Scr/Cond relative to Scr cells (Fig. 3C.iii, – lane 5 in Scr/Cond vs – lane 3 in Scr), and activation of downstream MAPK pathways (Fig. 3C.iv, – lane 5 in Scr/Cond vs – lane 3 in Scr) and NF-kB (Fig. 3C.v, – lane 5 in Scr/Cond vs – lane 3 in Scr). This induced cell proliferation, as assessed by elevated PCNA protein levels (Fig. 3C.vi, – lane 5 in Scr/Cond vs – lane 3 in Scr) and MTT assay (Fig. 3B.v, grey vs white bar). Additionally, *PPARγ1* mRNA levels were lowered in Scr/Cond-LX2 cells (Fig. 3B.iii, grey vs white bar), as expected during HSC activation and in contrast to the rise in *PPARβ/δ* levels [35], which was likely activated by the excess FAs produced in KD-LX2 and Src/Cond-LX2 cells (Table S2). Consistent with increased myofibroblastic transformation, *ACTA2* mRNA levels were induced by ∼two-to-threefold in KD and Scr/Cond LX2 cells (Fig. 3B.iv, grey and black vs white bar). Reversal of these processes in Scr/Cond cells by NA treatment (+ vs – lanes in Scr/Cond) further supported a role for FAs release from activated KD-LX2 in the autocrine activation of EGFR-Shc-MAPK to increase HSCs proliferation and activation.

Interleukin-6 (IL-6), a transcriptional target of NF-κB, was also elevated in KD media (Fig. 3Aii, black vs white bar). Consistent with the anti-inflammatory effect of NA [36] and its inhibition of IL-6 production [37], NA treatment reversed IL-6 level in KD media (Fig. 3Aii, + vs – lane) without affecting that of TNFα (Fig. 3Aiii, + vs – lane). Because IL-6 transactivates EGFR [38], we then tested whether the rise in IL-6 contributed to EGFR basal activation in KD-LX2 and Scr/Cond-LX2 cells. To this aim, we carried out media transfer experiments in the absence and presence of Gefitinib, an EGFR tyrosine kinase inhibitor [23]. As Fig. 4A shows, Gefitinib inhibited EGFR phosphorylation in KD-LX2 and Scr/Cond-LX2 cells (+ vs – lanes). In parallel, it reduced *PPARβ/δ (*Fig. 4B) and reciprocally induced *PPARγ1* mRNA levels in these cells (Fig. 4C, horizontally-hatched vs grey bar in Scr/Cond-LX2 and diagonally-hatched vs black bar in KD-LX2 cells) to stimulate their *CEACAM1* mRNA levels (Fig. 4D). This was associated with the ability of Gefitinib to prevent Scr/Cond-LX2 activation, as demonstrated by reduction and normalization of *ACTA2* mRNA levels (Fig. 4E) and their cell proliferation (Fig. 4F, + vs – lanes).

**Fig. 4.**
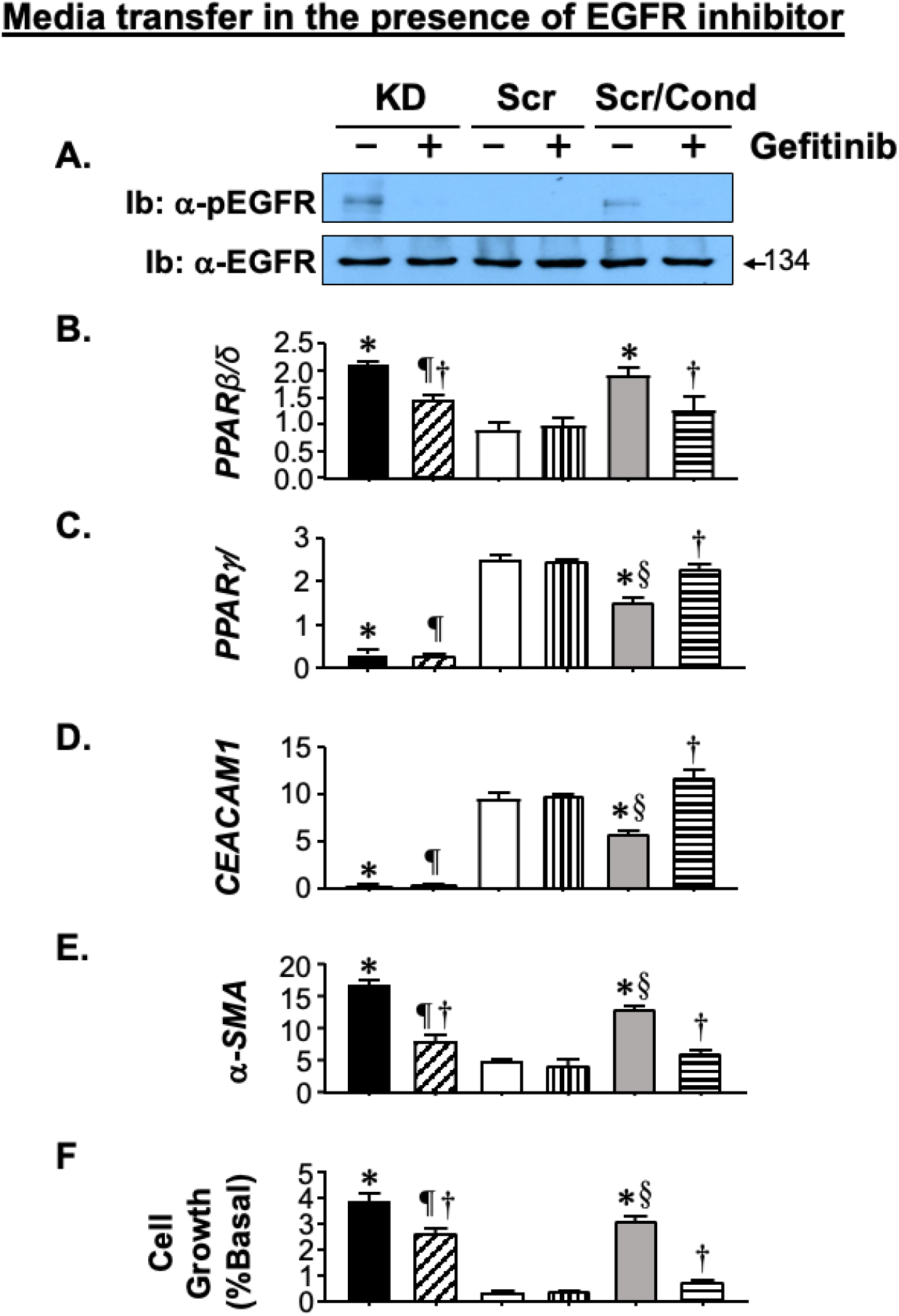
EGFR-mediated myofibroblastic transformation of KD-LX2 cells. Src-LX2 and KD-LX2 cells were incubated with Genitinib (10µM) or vehicle for 24 hrs. The conditioned media was collected from both Genitinib-treated and untreated KD-LX2 cells, and the supernatant was collected and transferred to Src-LX2 and incubated for 24 hrs (Scr/Cond-LX2 cells). (A) lysates were analyzed by immunoblotting (Ib) with α-pEGFR and α-EGFR for normalization to assess EGFR activation. Gel is representative of 2 different experiments performed on 2 different sets of mice. (B-E) lysates were analyzed by qRT-PCR analysis in triplicate to assess mRNA of *PPARβ/δ*, *PPARγ1*, *CC1,* and *ACTA2*. (F) cells were grown in 96 well plates to examine cell growth in triplicate by MTT assay (repeated twice). Values are expressed as mean ± SEM; \**P*<0.05 untreated KD and Scr/Cond vs untreated Scr-LX2 cells; ^†^*P*<0.05 NA-treated vs untreated/cell line; ^§^*P*<0.05 untreated Scr/Cond vs untreated KD, and ^¶^*P*<0.05 NA-treated KD vs NA-treated Scr and NA-treated Scr/Cond cells.

### 3.4. Activation of primary HSCs from Cc1^−/−^ null mice

In support of HSCs activation when their CEACAM1 is absent, primary HSCs from global *Cc1^−/−^* nulls exhibited higher mRNA levels of *Pparβ/δ*, *Pcna*, and *Acta2* than HSCs from wild-type mice (Table S3). They also exhibited higher *Srebp-1c* and *Fasn* mRNA levels. Moreover, their media induced the mRNA levels of these genes in wild-type HSCs (Table S3). NA treatment normalized these parameters in HSCs from *Cc1^−/−^* and *Cc1^+/+^/Cond* cells (Table S3). This proposed that CEACAM1 loss in HSCs activated them and caused their myofibroblastic transformation to contribute to hepatic fibrosis in *Cc1^−/−^* nulls [14].

### 3.5. *LratCre+Cc1^fl/fl^* mice with conditional deletion of *Ceacam1* in HSCs are insulin sensitive

Because *Ceacam1* loss in endothelial cells and hepatocytes could also contribute to hepatic fibrosis in *Cc1^−/−^* nulls [14], we then assessed the effect of deleting *Ceacam1* exclusively from HSCs on hepatic fibrosis. To this end, we generated *LratCre+Cc1^fl/fl^* mice with conditional deletion of *Ceacam1* in HSCs, as demonstrated by their intact *Ceacam1* expression in bone marrow macrophages, hepatocytes and endothelial cells (liver and heart) (Fig. 5A). These mice exhibited normal body weight, visceral fat mass, plasma NEFA and triacylglycerol levels (Table 1). *LratCre+Cc1^fl/fl^*mice exhibited normal hepatic insulin clearance (steady-state C-peptide/insulin molar ratio) and normo-insulinemia relative to their control counterparts (Table 1). They also showed normal tolerance to exogenous glucose and insulin (Fig. 5B and 5C, respectively), with normal fasting and fed blood glucose levels (Table 1). Consistent with normo-insulinemia, hepatic triacylglycerol levels were normal (Table 1) and H&E stain did not detect lipid droplet deposition in liver sections (Fig. 5D.d). Moreover, the mRNA levels of genes involved in fatty acid transport (CD36 translocase) and lipogenesis [Srebp-1c, and fatty acid synthase (Fasn)] were normal (Table S4). Together, this demonstrated that conditional *Ceacam1* deletion from HSCs did not cause insulin resistance or hepatic steatosis, consistent with intact expression of CEACAM1 in hepatocytes.

**Fig. 5.**
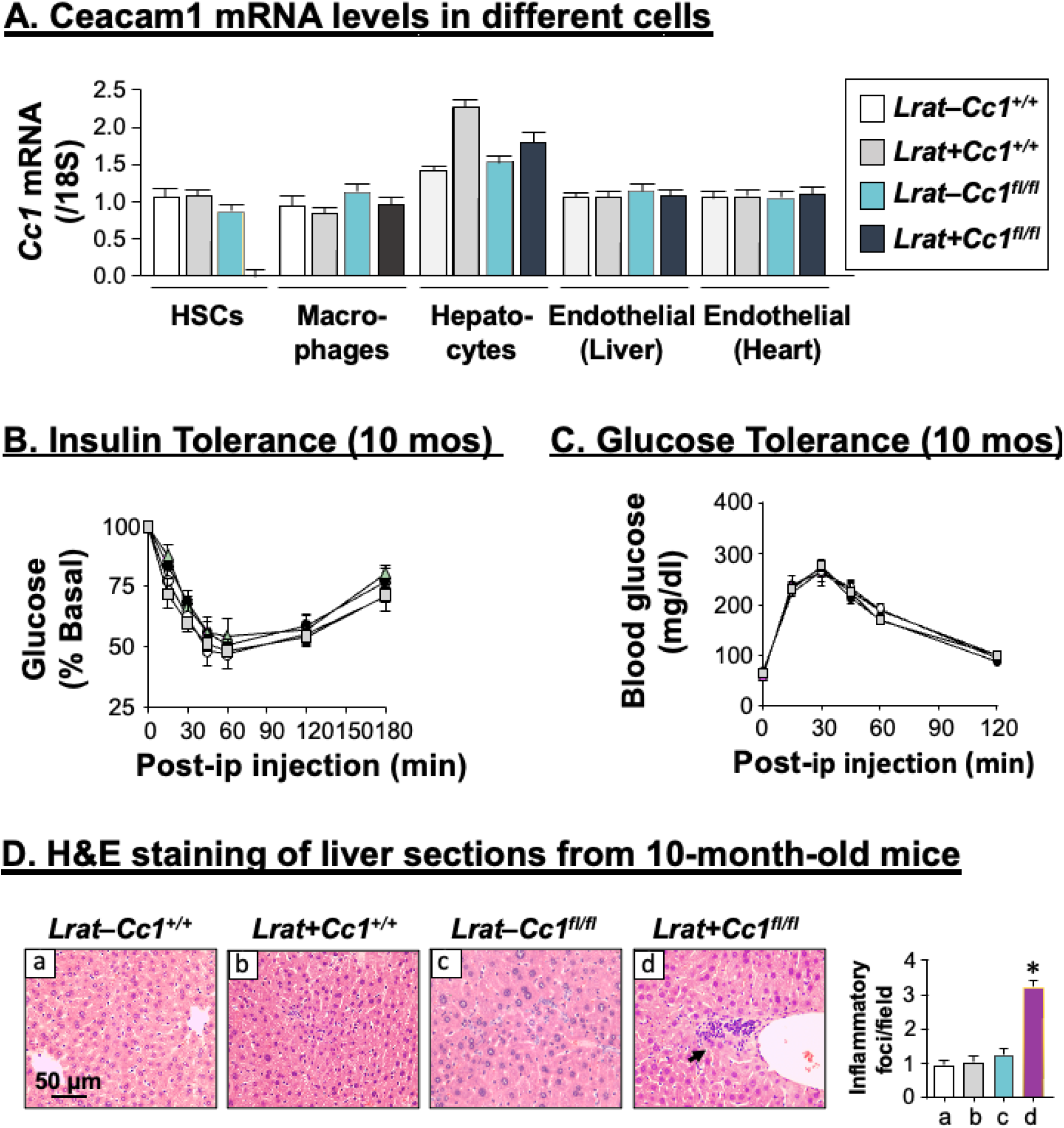
Metabolic phenotyping of *LratCre+Cc1^fl/fl^* mice. (A) Primary cells were isolated from male mutants and their littermate controls at 2-4 months of age except for HSCs which were isolated from mice at 8 months of age (n=2-5/genotype). *Ceacam1* mRNA levels were analyzed by qRT-PCR in triplicate and normalized to *18*s. Values are expressed as mean ± SEM. (B-C) 10-month-old male mice (n≥ 7–8/genotype) were injected intraperitoneally with insulin or glucose to assess glucose disposal in response to insulin (B) and glucose (C). Values were expressed as mean ± SEM. (D) Livers were removed from 10-month-old *Lrat+Cc^1fl/fl^* male mice and their 3 littermate controls (n=4–5/genotype), sectioned and stained with H&E staining to identify foci of inflammatory cell infiltrates in mutants (panel d) and their littermate controls (panels a-c). Values are expressed as mean ± SEM in the accompanying inflammatory foci quantification graph. \**P*<0.05 mutants vs the 3 littermate controls.

**Table 1.**
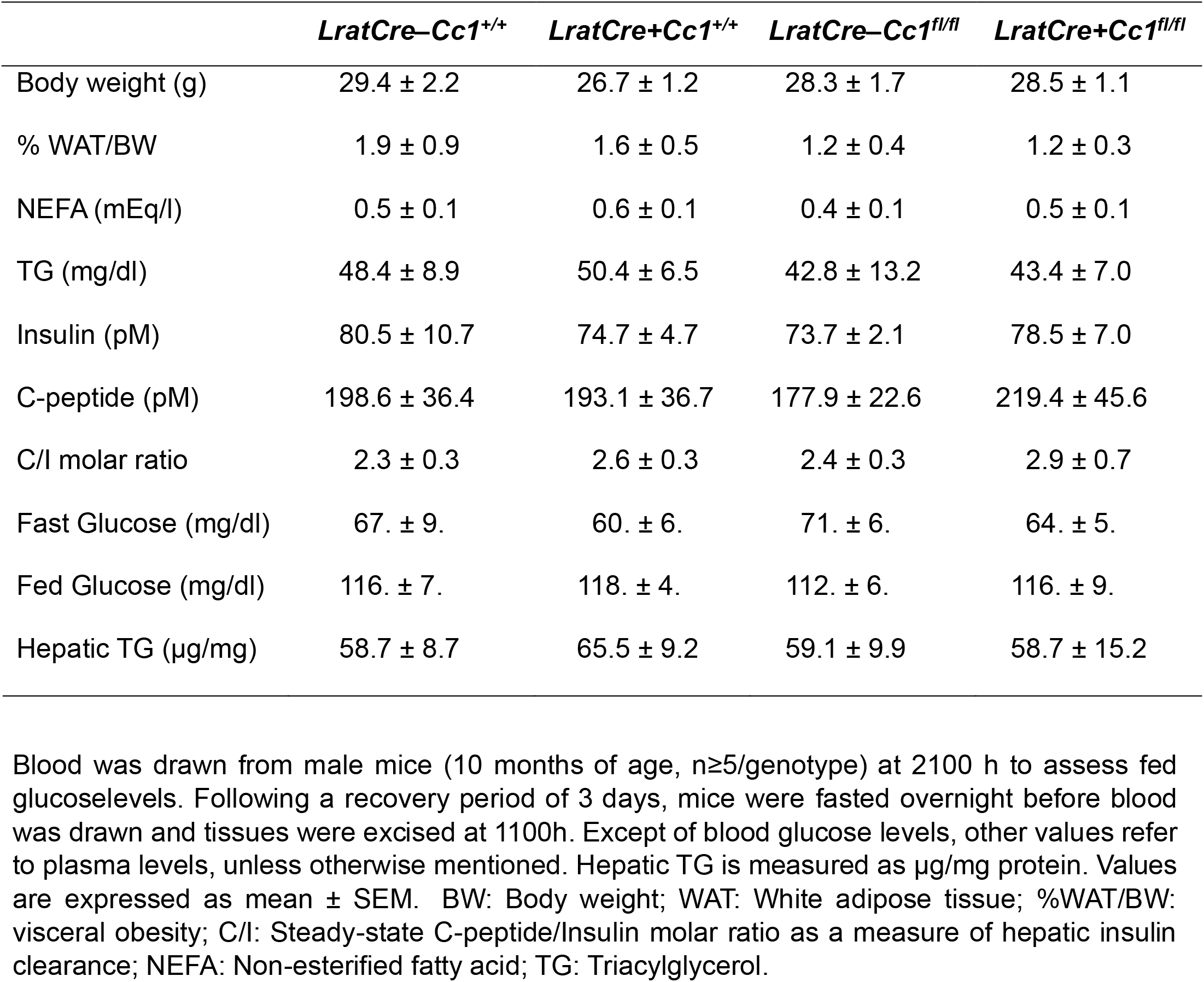
Plasma and tissue biochemistry in mice at 10 months of age.

|  | *LratCre–Cc1<sup>+/+</sup>* | *LratCre+Cc1<sup>+/+</sup>* | *LratCre–Cc1<sup>fl/fl</sup>* | *LratCre+Cc1<sup>fl/fl</sup>* |
| --- | --- | --- | --- | --- |
| Body weight (g) | 29.4 ± 2.2 | 26.7 ± 1.2 | 28.3 ± 1.7 | 28.5 ± 1.1 |
| % WAT/BW | 1.9 ± 0.9 | 1.6 ± 0.5 | 1.2 ± 0.4 | 1.2 ± 0.3 |
| NEFA (mEq/l) | 0.5 ± 0.1 | 0.6 ± 0.1 | 0.4 ± 0.1 | 0.5 ± 0.1 |
| TG (mg/dl) | 48.4 ± 8.9 | 50.4 ± 6.5 | 42.8 ± 13.2 | 43.4 ± 7.0 |
| Insulin (pM) | 80.5 ± 10.7 | 74.7 ± 4.7 | 73.7 ± 2.1 | 78.5 ± 7.0 |
| C-peptide (pM) | 198.6 ± 36.4 | 193.1 ± 36.7 | 177.9 ± 22.6 | 219.4 ± 45.6 |
| C/I molar ratio | 2.3 ± 0.3 | 2.6 ± 0.3 | 2.4 ± 0.3 | 2.9 ± 0.7 |
| Fast Glucose (mg/dl) | 67. ± 9. | 60. ± 6. | 71. ± 6. | 64. ± 5. |
| Fed Glucose (mg/dl) | 116. ± 7. | 118. ± 4. | 112. ± 6. | 116. ± 9. |
| Hepatic TG (µg/mg) | 58.7 ± 8.7 | 65.5 ± 9.2 | 59.1 ± 9.9 | 58.7 ± 15.2 |

Blood was drawn from male mice (10 months of age, n≥5/genotype) at 2100 h to assess fed glucose levels. Following a recovery period of 3 days, mice were fasted overnight before blood was drawn and tissues were excised at 1100h. Except of blood glucose levels, other values refer to plasma levels, unless otherwise mentioned. Hepatic TG is measured as µg/mg protein. Values are expressed as mean ± SEM. BW: Body weight; WAT: White adipose tissue; %WAT/BW: visceral obesity; C/I: Steady-state C-peptide/Insulin molar ratio as a measure of hepatic insulin clearance; NEFA: Non-esterified fatty acid; TG: Triacylglycerol.

### 3.6. Increased inflammation in LratCre+Cc1^fl/fl^ livers

H&E staining indicated diffused mononuclear inflammatory foci in the liver parenchyma of *LratCre+Cc1^fl/fl^* mutants without ballooning or altered hepatocellular architecture starting at 10 months of age [Fig. 5D.d (and graph) vs Fig. S3A.d from 8-month-old mice].

Immunohistochemical (IHC) analysis revealed increased macrophage recruitment (CD68) and activation (Mac2) [Fig. 6A.i-ii (and graphs), panels d vs a-c, respectively]. It also showed elevated immunostained myeloperoxidase (MPO) levels [Fig. 6A.iii (and graph), panel d vs a-c], which together with increased mRNA of MPO and elastase (Table S4), demonstrated an increase in neutrophil accumulation in the liver parenchyma of mutant livers. In addition to MPO, a granulocyte-specific transcription factor (STAT3) was activated (phosphorylated) at 10 months (Fig. 6B), but not at 8 months of age (Fig. S3B). This likely resulted from the ∼2-to-3-fold concomitant rise in hepatic IL-6 production (Fig. 6C vs Fig. S3C) and in the plasma levels of this pro-inflammatory cytokine (Fig. 6D).

**Fig. 6.**
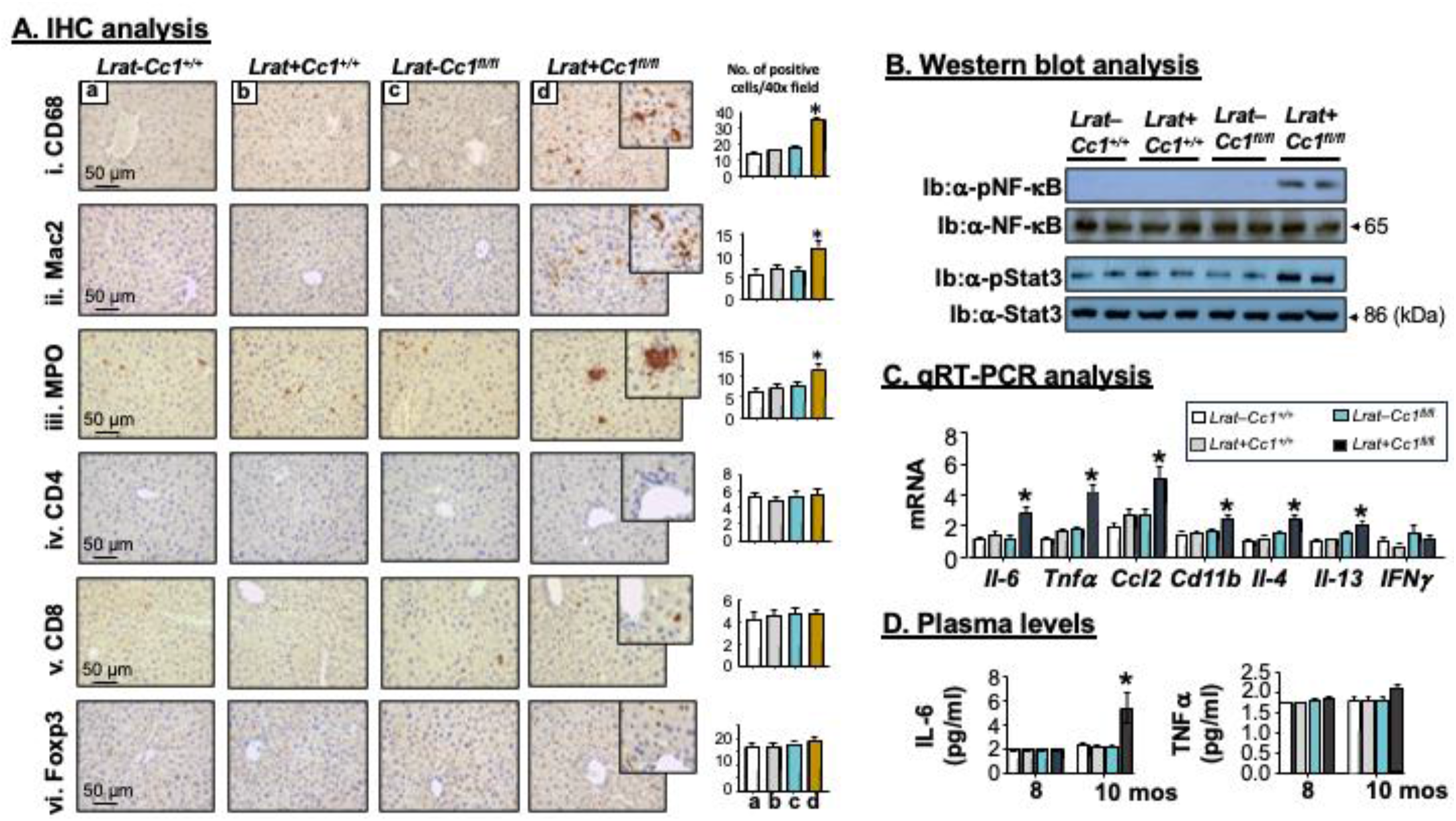
Increased inflammation in *LratCre+Cc1^fl/fl^* livers. (A) livers were removed from 10-month-old *LratCre+Cc1^fl/fl^* mutants and their three controls (n=4-5 mice/genotype) and subjected to (A) immunohistochemical (IHC) analysis with: (i) CD68 to assess macrophage recruitment, (ii) Mac-2 to examine macrophage activation, (iii) MPO to evaluate neutrophil accumulation, CD4 (iv) and CD8 (v) to immunostain T cells and (vi) Foxp3 to determine the anti-inflammatory Treg pool. Representative images taken at 50 μm magnification are shown with insets at 20 μm. Values are expressed as mean ± SEM in the accompanying quantification graph. \**P*<0.05 mutants (d) vs the 3 littermate controls (a-c). (B) liver lysates were subjected to immunoblotting (Ib) with antibodies against the phosphorylated p65 subunit of NF-κB (α-pNF-kB), and α-pStat3. To normalize against added proteins, gels were analyzed by SDS gel electrophoresis in parallel and proteins immunoblotted with specific antibodies. Representative gels include 2 different mice/genotype. (C) liver lysates (n=6/each genotype) were analyzed in duplicate by qRT-PCR using gene-specific primers and normalized to *Gapdh*. Values are expressed as mean ± SEM. \**P*<0.05 vs all three controls. (D) Male mice (8 and 10 months of age, n≥6/genotype/age group) were fasted overnight before blood was drawn at 1100 in the next morning and their plasma IL-6 and TNFα levels were analyzed. Values are expressed as mean ± SEM. \**P*<0.05 vs all three controls.

Consistent with IL-6 as a transcriptional target of NF-κB, the p65(NF-κB) subunit was basally activated (phosphorylated) in the livers of 10-month-old (Fig. 6B), but not 8-month-old (Fig. S3B) *LratCre+Cc1^fl/fl^* mutants. This likely resulted from reduced Shc sequestration in the absence of *Ceacam1* and the reciprocal increase in its coupling to EGFR [33]. In addition to IL-6, activated p65(NF-κB) could induce Mcp-1/Ccl2 transcription [39], as shown in Fig. 6C, to recruit monocytes/macrophages into active inflammatory foci in mutant livers. Together with IL-6, Ccl2 could induce CD11b+ macrophage pool (Fig. 6C) and its differentiation toward the M2 type [40; 41], which is partly mediated by elevated levels of IL-4/IL-13 type 2 cytokines (Fig. 6C). Together with no increase in the mRNA levels of Th1-derived cytokine IFNγ (Fig. 6C) or in plasma TNFα levels (Fig. 6D), this points to the mounting of an M2 response in mutant livers, mediated partly by sustained IL-6/STAT3 phosphorylation [42]. In contrast to macrophages, IHC (Fig. 6A.iv-v) and qRT-PCR (Table S4) analyses revealed no significant increase in pro-inflammatory CD4+T and CD8+T lymphocytes. Moreover, there was no increase in the anti-inflammatory Treg immunostain (Foxp3) (Fig. 6A.vi), or in hepatic IL-10 expression (Table S4). Thus, liver injury in *LratCre+Cc1^fl/fl^* mice was associated with a Th2 response marked by elevated IL-4/IL-13 secretion by hepatic lymphocytes that could activate infiltrated myeloid cells (macrophages and neutrophils) to induce their M2 genes expression.

### 3.7. Spontaneous fibrosis in LratCre+Cc1^fl/fl^ livers

Because activated hepatic macrophages could initiate and maintain the myofibroblastic transformation of HSCs [41], we then tested whether *LratCre+Cc1^fl/fl^* mice developed hepatic fibrosis. Based on Sirius Red staining, *LratCre+Cc1^fl/fl^*, but not their controls, developed an extensive interstitial chicken-wire pattern of collagen deposition starting at 10 months of age (Fig. 7A.d vs a-c, and vs Fig. S4A.d at 8 months). Consistently, the mRNA levels of pro-fibrogenic genes (*Acta2*, *Col1α1*, *Col3α1*, and *Tgfβ*) were induced in the livers of 10-month-old (Fig. 7B.i), but not 8-month-old (Fig. S4B) mutants. Hepatic fibrosis could be mediated by the activation of the canonical TGFβ–SMAD2/3 profibrogenic pathway, as demonstrated by SMAD2 phosphorylation (Fig. 7C) with no change in the expression of its inhibitor, Smad7 (Fig. 7B).

**Fig. 7.**
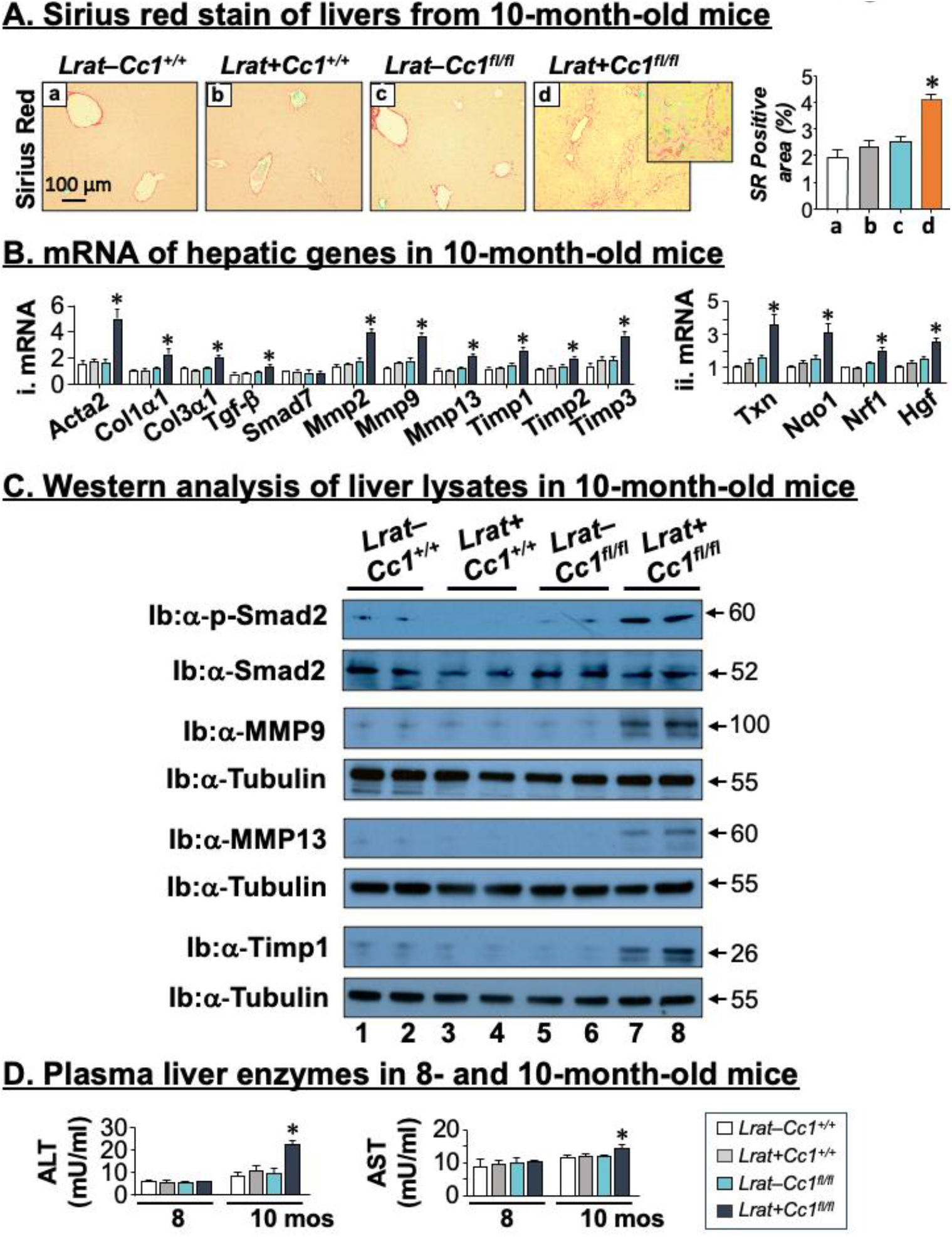
Spontaneous hepatic fibrosis in *LratCre+Cc1^fl/fl^* mice. Livers were removed from 10-month-old *Lrat+Cc^1fl/fl^* male mice and their 3 littermate controls (n=4–5/genotype). (A) Sirius red staining revealed increased deposition of interstitial chicken-wire pattern of collagen fibers in mutants (panel d) versus their littermate controls (panels a-c). Values are expressed as mean ± SEM in the accompanying quantification graph. \**P*<0.05 mutants vs the 3 littermate controls. (B) liver lysates (n=6/each genotype) were analyzed in duplicate by qRT-PCR using gene-specific primers and normalized to *Gapdh* to assess mRNA levels of genes involved in inflammation (i) and in hepatocytes injury (ii). Values are expressed as mean ± SEM. \**P*<0.05 vs all three controls. (C) Western Blot analysis of liver lysates from *LratCre+Cc1^fl/fl^* male mice (lanes 7-8) and their *LratCre–Cc1^+/+^* (lanes 1–2), *LratCre+Cc1^+/+^* (lanes 3-4) and *LratCre–Cc1^fl/fl^* (lanes 5– 6) controls. Phosphorylated Smad2 (α-pSmad2) normalized against α-Smad2. The protein levels of α-MMP9, α-MMP13 and α-Timp1 were normalized against α-Tubulin. Gels represent two different mice/genotype. The apparent molecular mass (kDa) is indicated at the right hand-side of each gel. (D) Male mice (8 and 10 months of age, n≥6/genotype/age group) were fasted overnight before blood was drawn at 1100 in the next morning and their plasma ALT and AST levels were analyzed. Values are expressed as mean ± SEM. \**P*<0.05 vs all three controls.

Activated HSCs modulate the extracellular matrix (ECM) composition, mediated by MAPK, NF-κB and TGFβ-SMAD2/3 pathways. This involves the regulation of the expression of the matrix metalloproteinases (MMPs) and the tissue inhibitor of metalloproteinases (TIMPs) that are implicated in the production as well as the resolution of excess collagen and other ECM components. Consistently, 10-month-old *LratCre+Cc1^fl/fl^*livers displayed higher mRNA (Fig. 7B.i) and protein levels (Fig. 7C) of MMP9, MMP13 and TIMP1 relative to controls. They also exhibited a ∼2-fold increase in the mRNA levels of hepatic *Mmp 2*, *Timp 2* and *Timp 3* (Fig. 7B.i). Whereas MMP9, TIMP1 and TIMP2 are pro-fibrogenic, MMP2 and TIMP3 block fibroblastic activation and increase collagen clearance [43]. MMP13 can promote collagen production as well as its clearance [43]. This demonstrated that the loss of CEACAM1 in HSCs regulated ECM formation by a complex of MMPs and TIMPs favoring hepatic fibrosis.

Coupled with oxidative stress, TGFβ signaling could cause hepatocellular injury [44]. Consistently, mutant livers manifested higher mRNA levels of genes implicated in oxidative stress (*Nox1* and *Nox4*) (Table S4) and hepatocytes injury (*Txn*, *Nqo*, *Nrf1* and *Hgf*) (Fig. 7B.ii). This could drive liver dysfunction, as determined by higher plasma alanine transaminase (ALT) and aspartate aminotransferase (AST) content in 10-month-old but not 8-month-old mutants as compared to control mice (Fig. 7D).

### 3.8. Conditioned media from LratCre+Cc1^fl/fl^ HSCs activates wild-type HSCs via an EGFR-mediated mechanism

As above, NA treatment blocked the release of FAs as glycerol (Fig. 8A.i, + vs – lane) and IL-6 (Fig. 8A.ii, + vs – lane) into the media of *LratCre+Cc1^fl/fl^* HSCs (KO). Thus, we next examined whether media from KO-HSCs (KO-Med) could activate wild-type (WT) HSCs and whether this could be blocked by NA treatment. As Fig. 8B.i shows, incubating WT-HSCs with KO-Med (WT/KO-Med) repressed *Ceacam1* expression by ∼65% relative to WT-HSCs incubated in regular culture media (Reg-Med) (– lanes, grey vs white bar). This likely resulted from increased *Pparβ/δ* and reduced *Pparγ_1_* expression (Fig. 8B.ii-iii, respectively, – lanes, grey vs white bars).

**Fig. 8.**
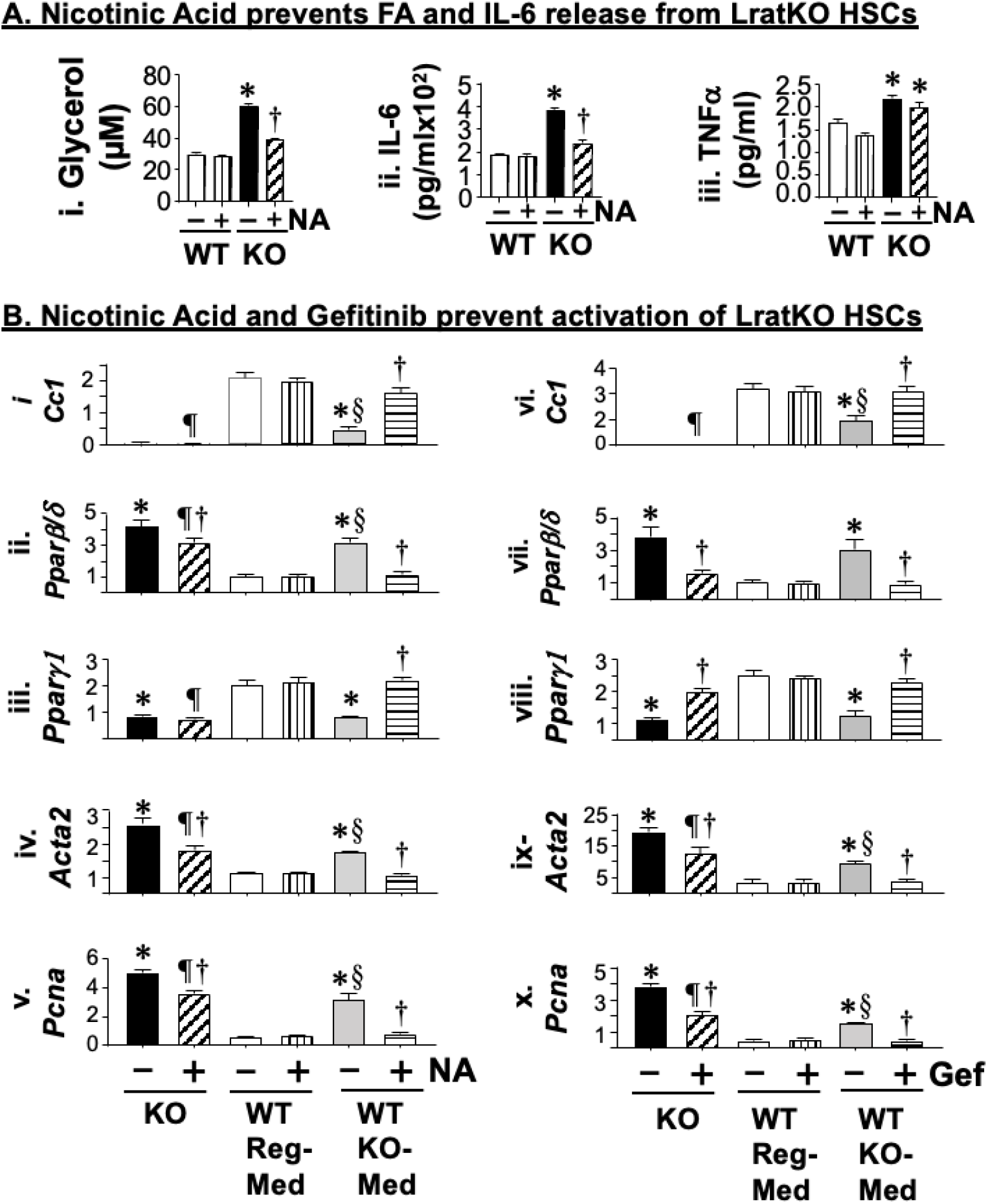
EGFR-mediated activation of *LratCre+Cc1^fl/fl^* HSCs. Primary HSCs were isolated from ≥8 *LratCre+Cc1^fl/fl^* mice (KO) and a combination of wild-type (*LratCre–Cc1^+/+^, LratCre+Cc1^+/+^* and *LratCre– Cc1^fl/fl^*) mice (WT). They were treated with (+) or without (–) NA and the media were collected and combined. (A) free glycerol (i), IL-6 (ii) and TNFα (iii) levels were assayed in the media. Values are mean ± SEM. *\*P*<0.05 KO (–) vs WT (–); ^†^*P*<0.05 NA-treated vs untreated/mouse group. (B) the conditioned KO-Med was transferred to WT HSCs (WT/KO-Med) for 24 hrs, while a parallel set of WT-HSCs was incubated in regular culture media (Reg-Med) and the mRNA levels were analyzed by qRT-PCR analysis. In some experiments, conditioned media without NA were transferred to WT-HSCs and the cells were treated with or without Gefinitib (vi-x). Cells were harvested for qRT-PCR analysis in triplicate of the mRNA levels of *Ceacam1* (i, vi), *Pparβ/δ* (ii, vii*), Pparγ1* (iii, viii)*, Acta2* (iv, ix) and *Pcna* (v, x). Values are expressed as mean ± SEM; \**P*<0.05 untreated KO and WT/KO-Med vs WT/Reg-Med; ^†^*P*<0.05 treated vs untreated/cell line; ^§^*P*<0.05 untreated WT/KO-Med vs untreated KO, and ^¶^*P*<0.05 treated KO vs treated WT/Reg-Med and treated WT/KO-Med cells.

This reciprocal change in *Pparβ/δ* and *Pparγ1* expression, together with higher expression of *Acta2* (Fig. 8B.iv) and *Pcna* (Fig. 8B.v) in WT/KO-Med than WT/Reg-Med cells (grey vs white bars) demonstrated a higher myofibroblastic activation and proliferation of WT/KO-Med than WT/Reg-Med cells. Furthermore, NA treatment reversed these changes in *Pparβ/δ*, *Pparγ1*, *Acta2* and *Pcna* mRNA levels in parallel to restoring *Ceacam1* expression in WT/KO-Med (Fig. 8B.i-vi, + vs – bars, horizontal vs vertical bars). Gefitinib had a similar effect on the mRNA of these genes in WT/KO-Med (Fig. 8B.vii-x, + vs – bars, horizontal vs vertical bars).

PPARβ/δ are activated by the FAs that are released from *LratCre+Cc1^fl/fl^* HSCs. As Table S5 shows, these KO-HSCs exhibited reduced RE synthesis with a reciprocal increase in PUFA-TG, as assessed by the 2-to-4–fold reduction in the mRNA levels of *Lrat* and *Lal/Lipa*, with the reciprocal ∼12-fold increase in Acsl4/Acsl1 and the ∼4-to-6–fold increase in the mRNA levels of *Dgat1* and *Atgl* (Table S5). It is likely that the increase in FAs release activated PPARβ/δ to reduce *Ceacam1* expression in WT/KO-Med. This would lower Shc sequestration and elevate its reciprocal coupling with EGFR to activate downstream pro-fibrogenic and proliferation pathways (increased *Acta2* and *Pcna*, respectively). Reversal of these changes in lipid metabolism in WT/KO-Med by Gefitinib further demonstrated that EGFR activation mediated the myofibroblastic transformation of HSCs by *Ceacam1* loss.

## 4. Discussion

The current study demonstrated that CEACAM1’s expression in cultured human LX2-HSCs is supported by autocrine PPARγ and retinoic acid transcriptional upregulation, and that activation of primary human HSCs significantly repressed CEACAM1 expression. On the other hand, loss of CEACAM1 in LX2 and primary murine HSCs activated them. This was manifested by reduced PPARγ_1_ and retinoic acid levels with reciprocal elevation in PPARβ/δ and PUFA-TG content, respectively. Because CEACAM1 inhibits FASN activity under normo-insulinemic conditions [45], suppressing *Ceacam1* transcription by PPARβ/δ [25] (and by the loss of PPARγ), likely mediated the increase in TG synthesis in mutant HSCs. In light of the anti-lipogenic and anti-fibrogenic effect of FASN inhibitors [46], the current data propose a key role for HSCs’ CEACAM1 in preventing hepatic fibrosis.

CEACAM1 expression is highest in hepatocytes. Its deletion in these cells impaired hepatic insulin clearance to cause hyperinsulinemia-driven insulin resistance, *de novo* lipogenesis and inflammation [15]. It also caused hepatic fibrosis, whereas hepatocytes-specific rescuing of CEACAM1 reversed steatosis and fibrosis in parallel to restoring insulin sensitivity in *Cc1^−/−^* null mice. This points to a key role for hyperinsulinemia-driven steatosis in hepatic fibrosis caused by CEACAM1 loss in hepatocytes [31]. In contrast, *Ceacam1* deletion from endothelial cells caused hepatic fibrosis in the absence of insulin resistance and hepatic steatosis [17]. The phenotype was driven by hyperactivation of the vascular endothelial growth factor (VEGFR)/NF-κB pathway and increased synthesis of endothelin1 and of its pro-fibrogenic signals via its receptor A in HSCs. Inflammation in this mutant preceded hepatic fibrosis and implicated macrophage activation in addition to mounting a Th1 response by T lymphocytes.

Like its deletion from endothelial cells, specific deletion of Ceacam1 from HSCs caused hepatic fibrosis in the absence of insulin resistance and hepatic steatosis. However, it occured concurrently to inflammation and was mediated by activation of EGFR by PUFA and IL-6, a transcriptional target of NF-kB. Sustained activation of the IL-6/STAT3 pathway could mediate the mounting of a Th2/M2 response in *LratCre+Cc1^fl/fl^* livers, as in *Stat1* nulls that exhibited activation of the M2 macrophage pool without a significant increase in pro-inflammatory T lymphocytes [47].

In addition to EGFR/NF-κB pathway, the EGFR/MAPK proliferative pathway was also activated in KD-LX2 HSCs devoid of CEACAM1. This resulted from the increased coupling of Shc to EGFR when its reciprocal sequestration by CEACAM1 was absent [33], as with respect to VEGFR [48] and the insulin receptor [32]. Thus, activation of NF-κB and the MAPK pathways downstream of these growth factor receptors constitutes a unifying mechanism underlying hepatic fibrosis when their shared substrate, CEACAM1, is lost. This agrees with the reported PPARβ/δ-driven HSCs proliferation and hepatic fibrosis via activation of the P38-JNK MAPK pathway in LX2 and murine HSCs [8].

EGFR is implicated in HSCs activation [34; 49] as demonstrated by the reversal of hepatic fibrogenesis, hepatocyte proliferation and liver injury in experimental models of hepatic fibrosis by inhibitors of EGFR tyrosine kinase activity [50; 51]. Yet, inhibiting EGFR phosphorylation to curb hepatic fibrosis has not gained traction at the clinical setting. Instead, targeting inflammation and lipogenesis constitutes the main current therapeutic approach, particularly at the early stages of the disease [2; 3]. This includes the use of a combinational therapy of PPARγ agonists and incretins to retard/attenuate hepatic fibrosis in patients with MASLD/MASH [52]. It is likely that the effectiveness of these drugs is mediated, at least partly, by the transcriptional activation of CEACAM1 [16], which would in turn, counter inflammation in immune cells [53] and lipogenesis in hepatocytes (by inhibiting FASN and limiting chronic hyperinsulinemia).

We have previously shown that hepatic CEACAM1 expression is progressively reduced with advancing fibrosis stages in patients with MASH [15]. Moreover, single cell RNA-sequencing showed lower CEACAM1 expression in hepatocytes and LSECs of patients with fibrosis/cirrhosis [17]. The current study demonstrated that activation of primary human HSCs repressed *CEACAM1* expression and that deleting *CEACAM1* from immortalized human LX2 activated them. The observations in *LratCre+Cc1^fl/fl^* mutants further emphasized the regulation of hepatic fibrosis by CEACAM1’s loss in HSCs, independently of its paracrine role in hepatocytes and endothelial cells. The underlying mechanisms converge at the level of NF-κB inflammation and MAPK proliferation pathways downstream of EGFR in HSCs (and in hepatocytes at the basal state) and of VEGFR in endothelial cells. Consistent with elevated serum IL-6 levels in patients with advanced hepatic fibrosis [54], loss of Ceacam1 in HSCs caused an elevation in plasma IL-6 levels concurrently wth hepatic fibrosis in *LratCre+Cc1^fl/fl^* mutants. Together, this proposes that inducing CEACAM1 expression could become an effective therapeutic approach to curb fibrosis, not only in early stages of the disease, but also at a later stage.

In summary, the current report provides an in vivo demonstration of a novel mechanistic link between a distinct CEACAM1/EGFR/NF-κB signaling module in murine HSCs and hepatic fibrosis, an advanced component of MASH. This was supported by studies in human LX2 HSCs demonstrating an autocrine regulation of hepatic fibrosis by the loss of CEACAM1 in stellate cells. Further analysis is required to translate our observations to MASH pathogenesis in humans.

## ABBREVIATIONS

CEACAM1: Carcinoembryonic Antigen-related Cell Adhesion Molecule 1 protein in mice and humans
*CEACAM1*: Human gene encoding CEACAM1 proteins
*Ceacam1*: Gene encoding CEACAM1 protein in mice
*LratCre+Cc1^fl/fl^*: Stellate cell-specific *Ceacam1* mutants
*LratCre–Cc1^+/+^*: Wild-type controls
*LratCre+Cc1^+/+^*: LratCre controls
*LratCre–Cc1^fl/fl^*: Ceacam1 Floxed controls
HSCs: Hepatic Stellate Cells
KD-LX2: LX2 human hepatic stellate cell line with shRNA-mediated suppression of CEACAM1
Scr-LX2: Control LX2 Line with scramble RNA.

## Credit authorship Contribution statement

H.T.M., H.E.G., S.A., S.G.L., S.V., H.L.S., S.Z., R.A., and G.D.B. researched data. H.T.M., H.E.G., and S.A.planned and organized experiments, collected and analyzed data, prepared the illustration and drafted the manuscript. S.L.F., R.F.S., L.A.vG, and T.D.H. Jr discussed data and edited the manuscript. S.M.N. conceived and oversaw the work, including its study design and data analysis, leading scientific discussions and reviewing/editing the manuscript.

## Declaration of competing interest

None declared.

## Acknowledgments

S.M.N. is partly supported by the Osteopathic Heritage Foundation J.J.Kopchick Eminent Research Chair.

## Financial support

This work was supported by NIH grants: R01-HL112248, R01-DK054254, R01-DK124126 (to S.M.N), R01-DK128289 (to S.L.F). S.V. is supported by FWO 1243121N, and L.A.vG. by FWO G071922N.

## Data availability statement

All data relevant to the study are included in the article/online supplemental information

## Supplemental Materials

### 1. Supplemental material and methods

#### 1.1. Luciferase assay

As described [1; 2], human hepatocellular carcinoma cells (HepG2) were cultured overnight to reach ∼60%-70% confluence. Transfection was performed with promoter constructs and the pRL-thymidine kinase *Renilla* luciferase promoter (Promega) using Lipofectamine 2000 (Invitrogen). Empty pGL4.10 vector was used as the negative control, pGL3-RARE-Luc (plasmid 13458; Addgene Cambridge, MA) and PPREx3-TK-Luc (Plasmid 1015; Addgene) were used as positive controls for PPRE and RXR, respectively. 24 hrs post-transfection, cells were serum-starved before being treated for 24 hrs with dimethyl sulfoxide (DMSO), 5 μM Retinoic Acid (Sigma-Aldrich Saint Louis, MO) and/or 1 μM Rosiglitazone (Sigma-Aldrich Saint Louis, MO). Luciferase activity was assessed using the Dual-Luciferase Reporter Assay System (Promega).

#### 1.2. Generation of human CEACAM1 shRNA lentiviral construct and treatment

The immortalized human hepatic stellate LX2 cells [initially characterized by the Friedman laboratory [3]] were cultured in Dulbecco’s Modified Eagle Medium (DMEM) High Glucose media (Gibco), supplemented with 2% FBS, 1% penicillin/streptomycin and 1% L-Glutamine. The free Web-based tool (http://www.genelink.com/sirna/shRNAi.asp) was used to design a putative siRNA against human CEACAM1 (hCEACAM1) and to design oligonucleotides that encode a corresponding small hairpin RNA (shRNA). Origene (Rockville, MD) constructed the shRNA plasmid with oligonucleotides: TGAATCCATGCCATTCAATGTTGCAGAGG and the homologous 120 sequence. The hCEACAM1 shRNA construct was co-transfected together with vectors expressing gag-pol, REV and VSV-G into 293FT cells (Invitrogen) to generate a third-generation lentiviral construct. Transfection was achieved by Lipofectamine 2000 (Invitrogen) using 100 ng total DNA per cm^2^ of the growth plate or well. The supernatants were harvested and the cell debris was removed by centrifugation at 2000g. The supernatant was used to infect human LX2 stellate cells after addition of 5 ng/ml polybrene (Sigma-Aldrich) and to establish ShCC1 line (KD) with stable downregulation of hCEACAM1 and scramble shRNA control (Scr). After 72 hrs, cells were selected by puromycin resistance (Gibco).

For insulin treatment, KD-LX2 and Scr-LX2 were serum-starved with phenol-free DMEM (Gibco) supplemented with 25 mM HEPES and 0.1%BSA for 16 hrs before being stimulated with or without 100 nM Insulin (Sigma-Aldrich) for 5 min before they were collected for Western blotting.

In some experiments, KD-LX2 and Scr-LX2 cells were stimulated with and without 100 nM Insulin (Sigma-Aldrich) for 24 hours and subjected to MTT assay (Sigma-Aldrich) and absorbance read at 570 nm in 96-well plates. Cell growth was calculated as percent of growth in the presence of insulin minus basal growth divided by maximum growth in complete medium [4].

#### 1.3. Generation of LratCre+Cc1^fl/fl^ mice

As in the conditional T cell-specific null mouse [5], the targeting Ceacam1 construct inserted a loxP-neo cassette in intron 6 and a loxP fragment in intron 9, deleting a sequence that encodes the cytoplasmic domain that is required for CEACAM1 phosphorylation by growth factor receptors [6; 7]. *Cc1loxp/loxp* mice were crossed with *LratCre* transgenic mice expressing a Cre recombinase directed by mouse lecithin-retinol acyltransferase [phosphatidylcholine-retinol-O-acyltransferase] (Lrat) promoter [8]. Heterozygotes were backcrossed >6x with C57BL/6J mice (Jackson laboratory). Stellate cell-specific deletion of *Ceacam1* in homozygotes (*LratCre+Cc1^fl/fl^* or *Lrat+Cc1^fl/fl^*), was confirmed by PCR using gene-specific primers (Fig. S1). As controls, we used homozygotes with wild-type *Ceacam1* allele with (*LratCre+Cc1^+/+^*; *Lrat+Cc1^+/+^*) or without Cre (*LratCre–Cc1^+/+^*; *Lrat–Cc1^+/+^*), and homozygotes with *Ceacam1*-floxed allele, without Cre (*LratCre–Cc1^fl/fl^*; *Lrat–Cc1^+/+^*). Genotyping of offspring was done by PCR analysis of ear DNA: FLOX A + FLOX B primers were used to detect the 383 bp wild-type allele (*Cc1^+/+^*), FLOX A + FLOX C detected the 488 bp knockout-type allele (*Cc1^fl/fl^*). The *LratCre* reaction detected a 343 bp band in the *LratCre+Cc1^+/+^* and *LratCre+Cc1^fl/fl^*.

#### 1.4. Glucose and insulin tolerance tests

As routinely done [9], male mice were kept in cages with Alpha-dri bedding (Shepherd Specialty Papers) and fasted for 6 hrs before being injected intraperitoneally with either 1.5g/kg BW dextrose solution (GTT or 0.75units/kg BW human regular insulin (Novo Nordisk, Princeton, NJ) (ITT) and their tail blood glucose was measured at 0-120 min (GTT) or 0-180 post-injection (ITT).

#### 1.5. Biochemical parameters

Following their move to Alpha-dri bedding (Shepherd Specialty Papers), mice were fasted overnight for 18 hrs, anesthetized with an IP injection of pentobarbital (1.1mg/kg BW) at 1100h the next morning and their venous blood drawn into heparinized micro-hematocrit capillary tubes (Catalog number 22-362566, Fisherbrand, Waltham, MA). Plasma was processed and tissues excised to determine hepatic triacylglycerol as previously described [10]. Plasma was analyzed for insulin (80-INSMSU-E01 ELISA kit; Alpco, Salem, NH), C-peptide (80-CPTMS-E01 ELISA kit; Alpco), non-esterified fatty acids (NEFA-C enzymatic colorimetric assay; Wako, Richmond, VA), Tumor necrosis factor-alpha (TNFα ELISA Kit, ab100747, Abcam), Interleukin-6 (IL-6 ELISA Kit, ab222503, Abcam), alanine transaminase (ALT ELISA KIT, 700260; Cayman Chemical Company, Ann Arbor, MI), and aspartate aminotransferase (AST ELISA, 701640; Cayman Chemical Company).

#### 1.6. Liver histology and immunohistochemical analysis

As routinely done, liver sections were fixed in 10% neutral buffered formalin (Sigma-Aldrich, Milwaukee, WI), paraffin-embedded, cut, mounted and stained with hematoxylin-eosin (H&E) to assess steatosis and lobular inflammation. Fibrosis was assessed by staining deparaffinized and rehydrated slides with 0.1% Sirius Red stain (Direct Red 80, Sigma-Aldrich, St Louis, MO) in saturated aqueous solution of picric acid (Sigma-Aldrich), washed in 0.5% acetic acid (Sigma-Aldrich), hydrated in different alcohol percentages, and cleared in Xylene (Sigma-Aldrich). Images were taken using Nikon Eclipse 90i Microscope (Nikon, Melville, NY) as described [11]. 10 randomly selected high power fields (20x) per sample were imaged and quantified with ImageJ (v1.53t) to determine the percentage of Sirius red (SR) stain area within the whole area of imaged hepatic tissue. Briefly, each image was RGB-stacked, subjected to standardized thresholding of individual channel to isolated Sirius Red stain and then quantified as % area. Image quantifications were averaged and the mean within each experimental group was plotted in the accompanying graph.

For immunohistochemical (IHC) analysis, FFPE sections were deparaffinized, dewaxed in xylene, rehydrated before quenching endogenous peroxidases with 3% H_2_O_2_ buffer. Antigen-retrieval was then performed by incubating slides in 10mM sodium citrate buffer (pH 6.0) or 10 mM Tris-EDTA buffer (pH 9.0) in a microwave. Blocking was done using 1% rabbit or mouse serum (Vector labs, Burlingame, CA) for 1 hr at room temperature. Slides were stained overnight at 4°C in a humidifier chamber with α-CD68 (1:150, polyclonal, Abcam, Cambridge, MA), α-Mac2 (1:250 monoclonal, Abcam), α-MPO (1:1000 monoclonal, Abcam),, α-CD4 (1:1000 monoclonal, Abcam), α-CD8 (1:2000 monoclonal, Abcam), and α-Foxp3 (1:100 Invitrogen, Waltham, MA). Slides were then incubated with appropriate species-specific biotinylated ImmPRESS HRP horse anti-mouse or anti-rabbit secondary antibody (Vector Labs, Burlingame, CA) for 30min before being treated with 3,3’-Diaminobenzidine (DAB, Vectastain kit-Vector labs, Newark, CA) and counterstain with Mayer’s hematoxylin. Images were taken using Nikon Eclipse 90i Microscope (Nikon, Melville, NY), as previously described [11]. Sections were evaluated blindly, and positively stained cells were counted in 5 fields/mouse at 40X magnification.

#### 1.7. Isolation of primary murine cells

Primary hepatocytes, bone marrow-derived macrophages, and endothelial cells from livers and hearts were isolated from ketamine/xylazine-anesthetized 2-to-3–month-old male mice, as previously described [11]. Briefly, hepatocytes [12] were maintained in DMEM (Gibco Lab, Gaithersburg, MD)-10% FBS, 1% glutamine, and 1% penicillin/streptomycin (Gibco). Bone marrow-derived macrophages [11] were obtained by flashing femurs and tibias with sterile Roswell Park Memorial Institute (RPMI)-1640 medium from Gibco supplemented with 10% horse serum, 1% L-glutamine, 1% Penicilin/Streptomycin and 10ng/ml Recombinant Mouse factor macrophage colony-stimulating factor [M-CSF] R&D Systems, Minneapolis, MN). Primary endothelial cells [13; 14] isolated from livers and hearts were maintained in DMEM-F12 (Gibco) supplemented with 20% FBS, 1% glutamine, and 1% penicillin/streptomycin (Gibco), 100 ug/ml Heparin (Sigma-Aldrich) and 100ug/ml Endothelial Cell Growth supplement [ECGS) (EMD Millipore Corporation Burlington MA).

Primary mouse hepatic stellate cells (HSCs) were isolated from ≥8-month-old mice, as described [15]. Mice were anesthetized and using laparotomy, liver and the inferior vena cava (IVC) were exposed and liver were perfused with *in situ* with EGTA solution (8,000 mg/l NaCl, 400 mg/l KCl, 88.17 mg/l NaH_2_PO_4_.H_2_O, 120.45 mg/l, NaHPO4 2,380 mg/l HEPES, 350 mg/l NaHCO3, 190 mg/l EGTA, and 900 mg/l Glucose) for 1-2 min, at a rate of 5ml/min. This was followed by perfusion with the Pronase solution for 5 min at 5ml/min. Pronase solution was prepared by dissolving 14 mg/mouse of pronase (Sigma-Aldrich) in 35 ml of enzyme buffer solution (8,000 mg/l NaCl, 400 mg/l KCl, 88.17 mg/l NaH_2_PO_4_, 120.45 mg/l, NaHPO_4_ 2,380 mg/l HEPES, 350 mg/l NaHCO_3_, and 560 CaCl2.2H_2_O). The livers were then perfused for 7 min with collagenase solution composed of 3.7 U/mouse of Collagenase D (Roche, Pleasanton, CA) dissolved in 40 ml enzyme buffer solution. After perfusion, the partially digested livers were excised, placed into a sterile Petri dish containing Pronase/Collagenase solution prepared by dissolving 25 mg/mouse of pronase and 4.4 U/mouse to 50 ml enzyme buffer solution and minced under cell culture hood. 1% DNase was added (Roche, Pleasanton, CA) before further digesting the liver at 40^0^C for 20 min and the digest passed through a 70m cell strainer (Sigma-Aldrich) to remove undigested materials. The cell suspension was centrifuged at 580g for 10 min at 4^0^. The pellet was then washed using 4°C Gey’s buffered salt solution B (GBSS/B) (8,000 mg/l NaCl, 370 mg/l KCl, 210 mg/l MgCl_2_ .6H_2_O, 70 mg/l MgSO_4_ 7H_2_O, 75 mg/l Na_2_HPO_4_ 2H_2_O, 30 mg/l KH_2_PO_4_, 991 mg/l glucose, 227 mg/l NaHCO_3_, and 225 mg/l CaCl_2_ H_2_O, supplemented with DNase I by centrifugation at 580g at 4^0^C for 10 min. HSCs were purified from the rest of cells including hepatocytes by density gradient-mediated separation. Cells were resuspended in 32 ml Gey’s buffered salt solution (GBSS) before 16 ml Nycodenz solution [4.94 g of Nycodenz (Accurate Chemicals Westbury, NY) in 15 ml GBSS/B without NaCl] was added. Gently overlaid cells-Nycondenz suspension with 1.5 ml GBSS using 3 ml syringe with a 26-gauge needle. Centrifuge the cell suspension at 1.380g at 4^0^C for 17 min without brake. The interphase containing enriched HSC between the GBSS and Nycondenz layer was removed and washed using GBSS by centrifugation at 580g, 4^0^C for 10 min. The cell pellet containing HSCs was resuspended in prewarmed culture media DMEM (Gibco, Gaithersburg, MD) supplemented with 10% fetal bovine serum (FBS), 0.04% Gentamycin, and 1% Antibiotic/antimyocotic (Gibco), placed in a humidified tissue culture incubator (37°C, 5%CO_2_).

#### 1.8. Western blot analysis

Proteins were extracted from tissues or cells using lysis buffer containing 150 mM NaCl-50 mM Hepes (pH 7.6), 0.02% sodium azide,1% Triton X-100, 50 mM NaF, PMSF, Na_3_VO_4_ and proteinase Inhibitor Cocktail Tablet (Roche Diagnostics Penzberg, Germany). Protein concentration was determined by Bio-Rad Protein Assay (Bio-Rad Laboratories, Hercules, CA) and 10-30 µg proteins were denatured by boiling in sodium dodecyl sulfate (SDS) buffer for 3 min and resolved by 7% SDS-PAGE, transferred onto nitrocellulose membrane (Bio-Rad Laboratories). After blocking with 3%-5% dry milk or bovine serum albumin (BSA) dissolved in TBST (Tris-Buffered Saline, pH 7.2, 0.1% Tween 20), the membrane was incubated at 4^0^C overnight with polyclonal antibodies (1:1000): phospho-Akt ^Ser^ ^473^, Akt, phospho-p44/42 MAPK ^(Thr^ ^202/Tyr204)^, p44/42 MAPK, phospho-SMAD2^Ser^ ^465/467^, SMAD2, phospho-SMAD3^Ser^ ^423/425^, SMAD3, phospho-EGFR1 ^(Y1173)^, EGFR, Shc, phospho-NF-κB p65 (Ser536), NF-κB, phospho-Stat3 (Tyr705) Stat3, MMP9, MMP13 and Timp1 (Cell signaling, Danvers, MA), phospho-Insulin receptor beta (pIR_β_) (phospho-Y1361), Insulin receptor beta (C18C4) (Abcam), PCNA (Santa Cruz, Dallas, TX) , phospho-tyrosine (4G10) (EDM Milipore, Billerica, MA). The following custom-made rabbit polyclonal antibodies were used: Ab 3759 against the mouse CEACAM1 extracellular domain, and anti-human CEACAM1 clone 18/20, a gift from the late Dr. Bernhard Singer (University Hospital Essen, Germany). Next day the membrane was washed three times with TBST and incubated with horseradish peroxidase conjugated secondary antibodies antibody (GE Healthcare Life Sciences, Amersham, Sunnyvale, CA, USA), for one hour at room temperature. For normalization, membranes were reprobed with monoclonal antibodies against tubulin (Cell signaling). Proteins were detected by chemiluminescence (Thermo Scientific, Rockford, IL).

| Name | Company | Catalog Number |
| --- | --- | --- |
| **Primary Antibodies for Western blot** |  |  |
| Akt | Cell Signaling Technology | 9272 |
| phospho-Akt (Ser473) | Cell Signaling Technology | 4051 |
| EGFR1 | Cell Signaling Technology | 4267 |
| phospho-EGFR1 (Y1173) | Cell Signaling Technology | 4407 |
| Insulin receptor beta | Abcam | ab69508 |
| phospho-Insulin receptor beta (pIR<sub>β</sub>)<br>(phospho-Y1361) | Abcam | Ab303492 |
| p44/42 MAPK | Cell Signaling Technology | 9102 |
| phospho-p44/42 MAPK (Thr202/Tyr204) | Cell Signaling Technology | 9101 |
| Phosphotyrosine (4G10) | EDM Millipore | 05-321 |
| MMP2 | Cell Signaling Technology | 40994 |
| MMP9 | Cell Signaling Technology | 13667 |
| MMP13 | Cell Signaling Technology | 69926 |
| NF-κB p65 | Cell Signaling Technology | 8242 |
| Phospho-NF-κB p65 (Ser536) | Cell Signaling Technology | 3031 |
| PCNA | Santa Cruz, Dallas | Sc-56 |
| Shc | Cell Signaling Technology | 50832 |
| Smad2 | Cell Signaling Technology | 5339 |
| phospho-Smad2 (Ser465/467) | Cell Signaling Technology | 3108 |
| Smad3 | Cell Signaling Technology | 9523 |
| phospho-Smad3 (Ser423/425) | Cell Signaling Technology | 9520 |
| Stat3 | Cell Signaling Technology | 1264 |
| phospho-Stat3 (Tyr705) | Cell Signaling Technology | 9145 |
| TIMP1 | Cell Signaling Technology | 8946 |
| Tubulin | Cell Signaling Technology | 2148 |
| **Secondary Antibodies for Western blot** |  |  |
| Donkey anti-rabbit IgG antibody (HRP) | GE Healthcare Life Sciences | NA934V |
| Sheep anti-mouse IgG (HRP) | GE Healthcare Life Sciences | NXA931V |
| **IHC Antibodies-Mouse** |  |  |
| CD4 | Abcam | ab183685 |
| CD8 | Abcam | ab209778 |
| CD68 | Abcam | ab125212 |
| Foxp3 | Invitrogen | 700914 |
| MPO | Abcam | ab208670 |
| Mac2 | Abcam | ab76245 |

---

**Table S1.**
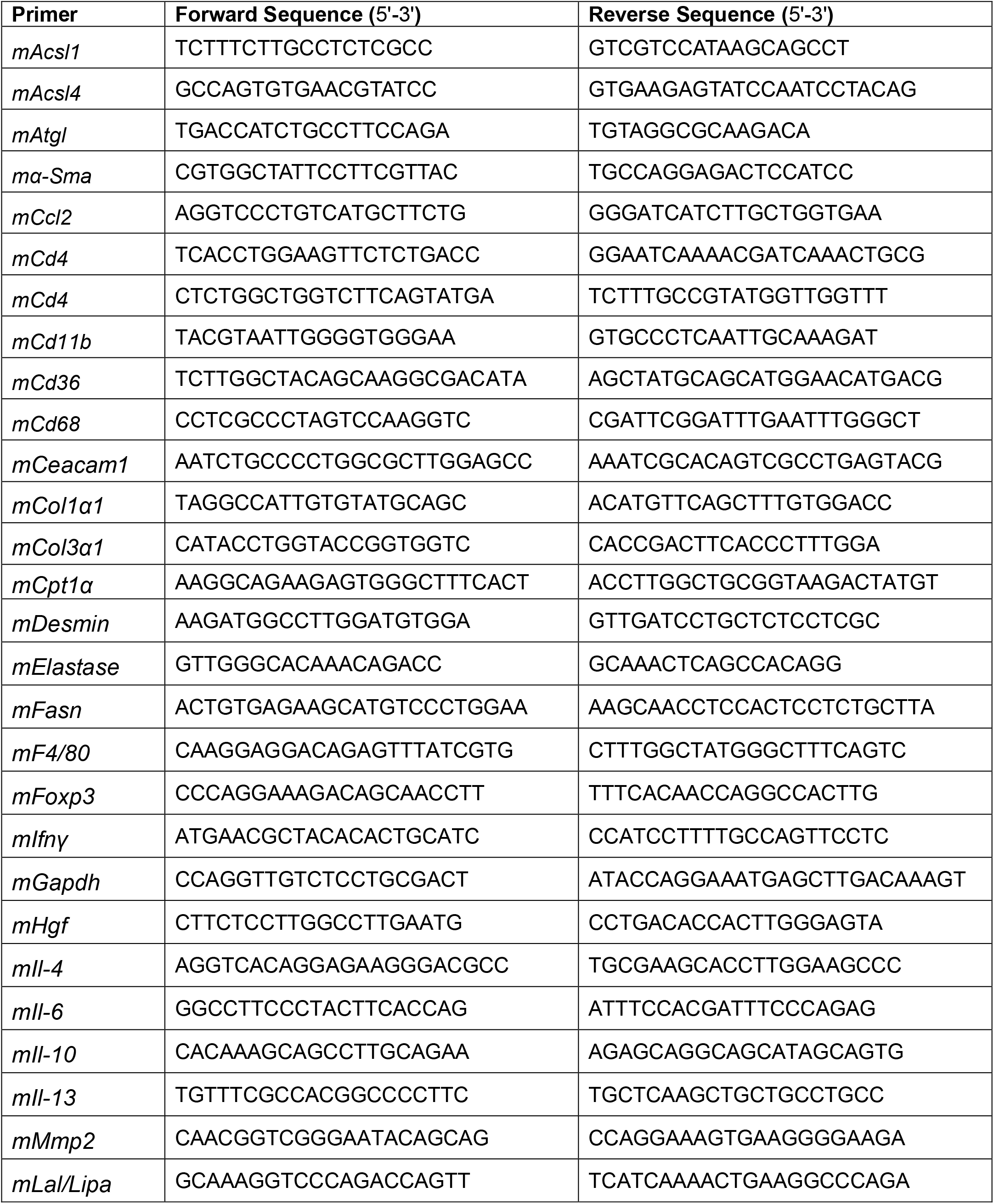
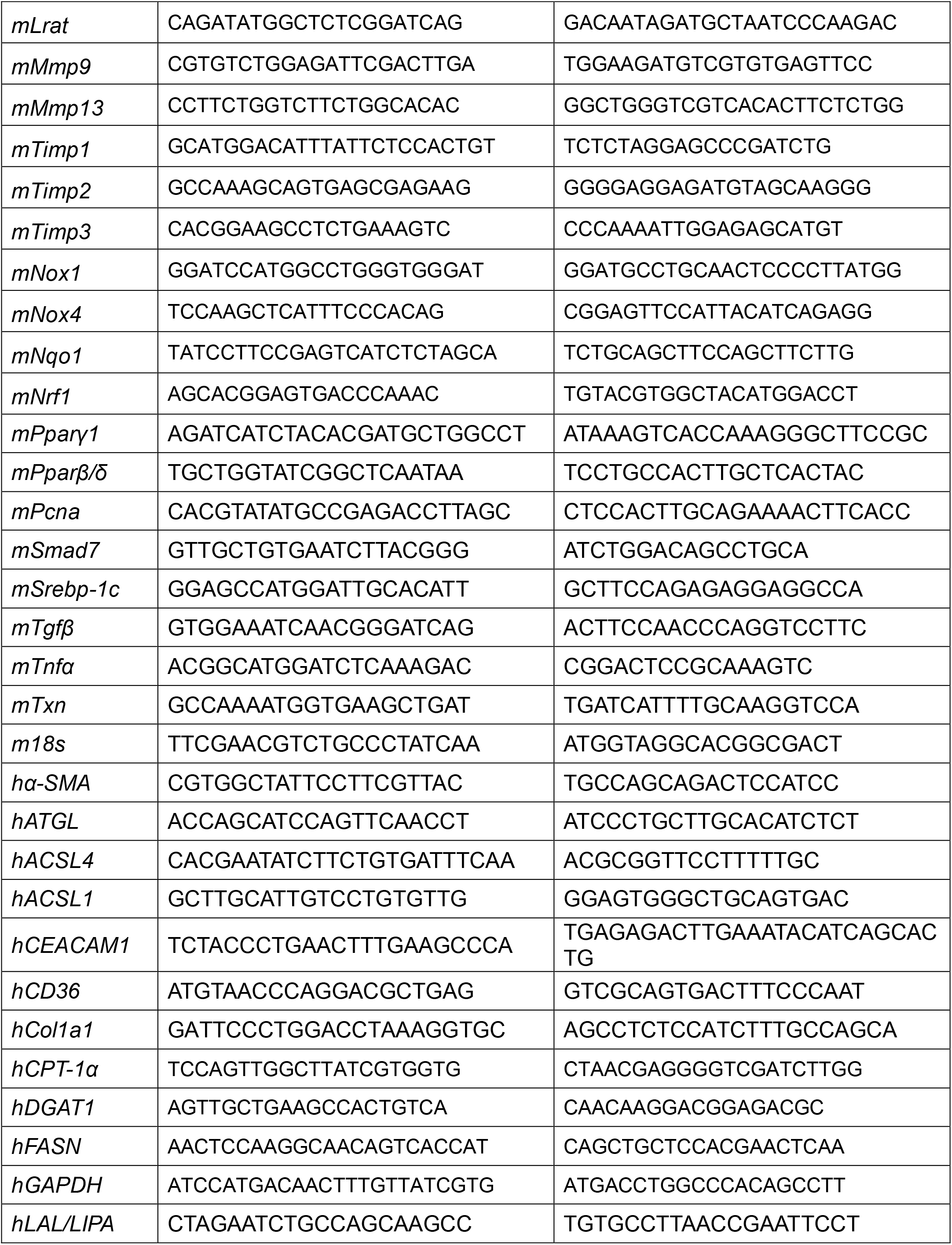
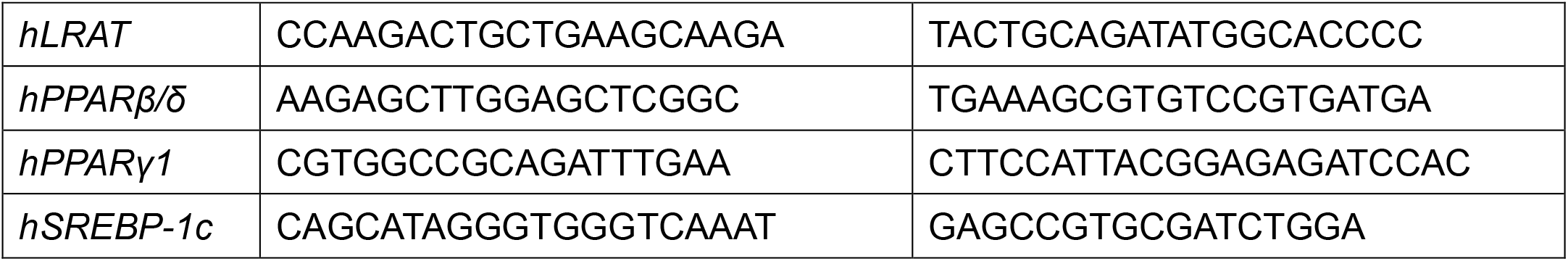
Primer sequences of mouse *(m)* and human *(h)* genes used in qRT-PCR.

**Table S2.**
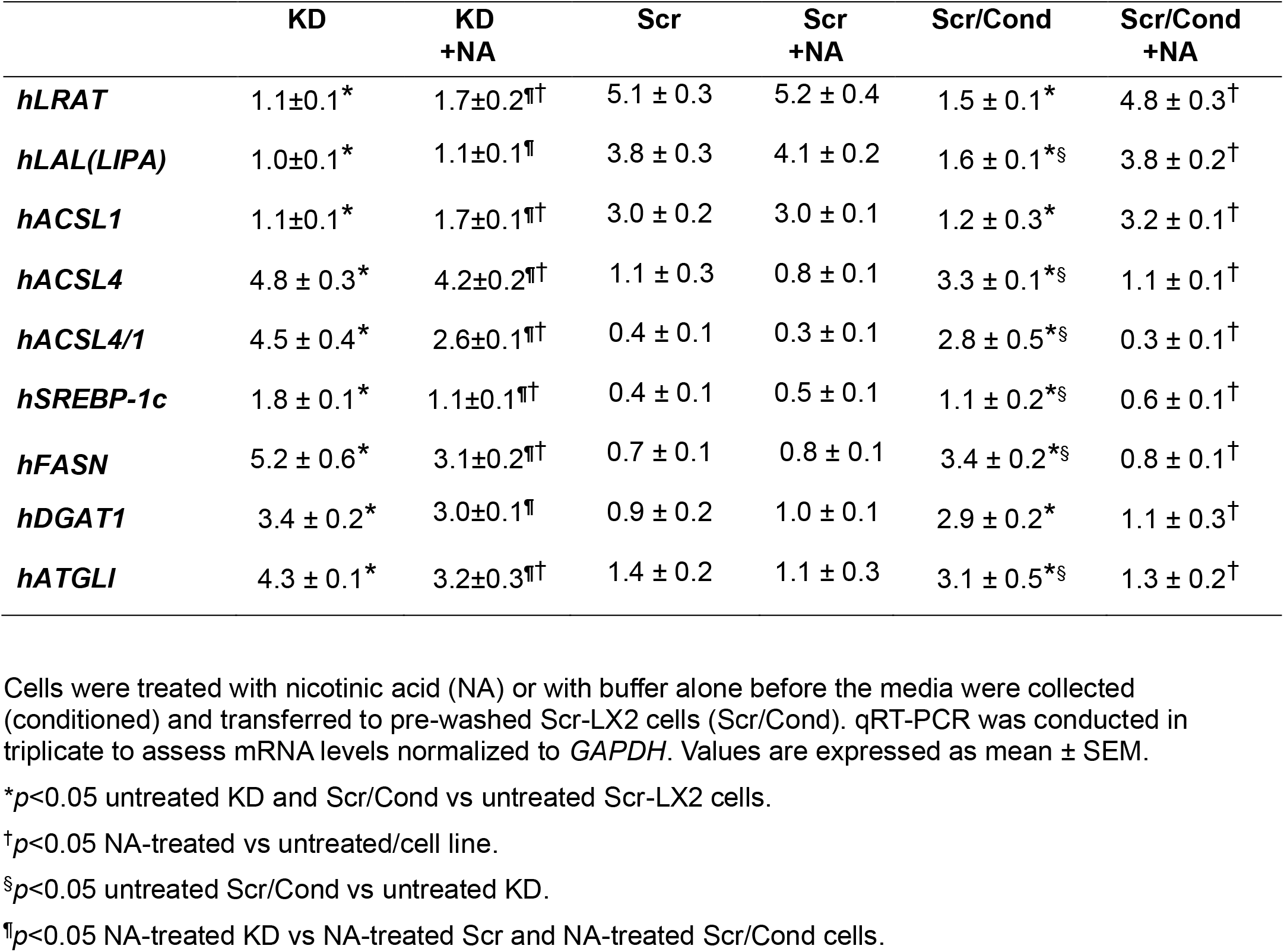
The effect of nicotinic acid on lipid metabolism of HSCs from KD-LX2 transferred into Scr-LX2 HSCs.

**Table S3.** qRT-PCR analysis in primary cells incubated in conditioned media from *Cc1^−/−^* HSCs.

|  | *Cc1<sup>-/-</sup>* | *Cc1<sup>-/-</sup>*<br>+NA | *Cc1<sup>+/+</sup>* | *Cc1<sup>+/+</sup>*<br>+NA | *Cc1<sup>+/+</sup>/Cond* | *Cc1<sup>+/+</sup>/Cond* +NA |
| --- | --- | --- | --- | --- | --- | --- |
| ***Ceacam1*** | negligible | negligible | 1.12±0.01 | 1.05±0.02 | 0.53±0.02*§ | 1.08±0.03<sup>†</sup> |
| ***Pparβ/δ*** | 2.12±0.01* | 1.45±0.03<sup>††</sup> | 1.11±0.01 | 1.05±0.02 | 1.94±0.03* | 1.05±0.03<sup>†</sup> |
| ***Pcna*** | 2.27±0.02* | 1.43±0.03<sup>††</sup> | 1.07±0.04 | 1.02±0.02 | 2.03±0.03* | 1.06±0.03<sup>†</sup> |
| ***α-Sma*** | 2.29±0.02* | 1.44±0.03<sup>††</sup> | 1.12±0.01 | 1.05±0.02 | 2.13±0.04* | 1.08±0.03<sup>†</sup> |
| ***Srebp-1c*** | 1.28±0.03* | 0.58±0.02<sup>††</sup> | 0.34±0.02 | 0.34±0.02 | 0.83±0.02* | 0.36±0.02<sup>†</sup> |
| ***Fasn*** | 2.38±0.01* | 1.64±0.02<sup>††</sup> | 1.25±0.03 | 1.26±0.03 | 2.01±0.03* | 1.40±0.06<sup>†</sup> |

Primary HSCs were isolated from ≥8-month-old mice (n=7/genotype), pre-incubated with (+) or without (-) nicotinic acid (NA) before *Cc1<sup>-/-</sup>* media were collected and transferred into *Cc1<sup>+/+</sup>* control HSCs (*Cc1<sup>+/+</sup>/cond*). qRT-PCR analysis was carried out in triplicate, normalized against *Gapdh* and presented as mean ± SEM.
\**p*<0.05 untreated HSCs from *Cc1<sup>-/-</sup>* and *Cc1<sup>+/+</sup>/Cond* vs untreated *Cc1<sup>+/+</sup>* cells.
<sup>†</sup>*p*<0.05 NA-treated vs untreated/cell line.
§*p*<0.05 untreated *Cc1<sup>+/+</sup>/Cond* vs untreated *Cc1<sup>-/-</sup>*.
<sup>††</sup>*p*<0.05 NA-treated *Cc1<sup>-/-</sup>* vs NA-treated *Cc1<sup>+/+</sup>* and NA-treated *Cc1<sup>+/+</sup>/Cond* cells.

**Table S4.**
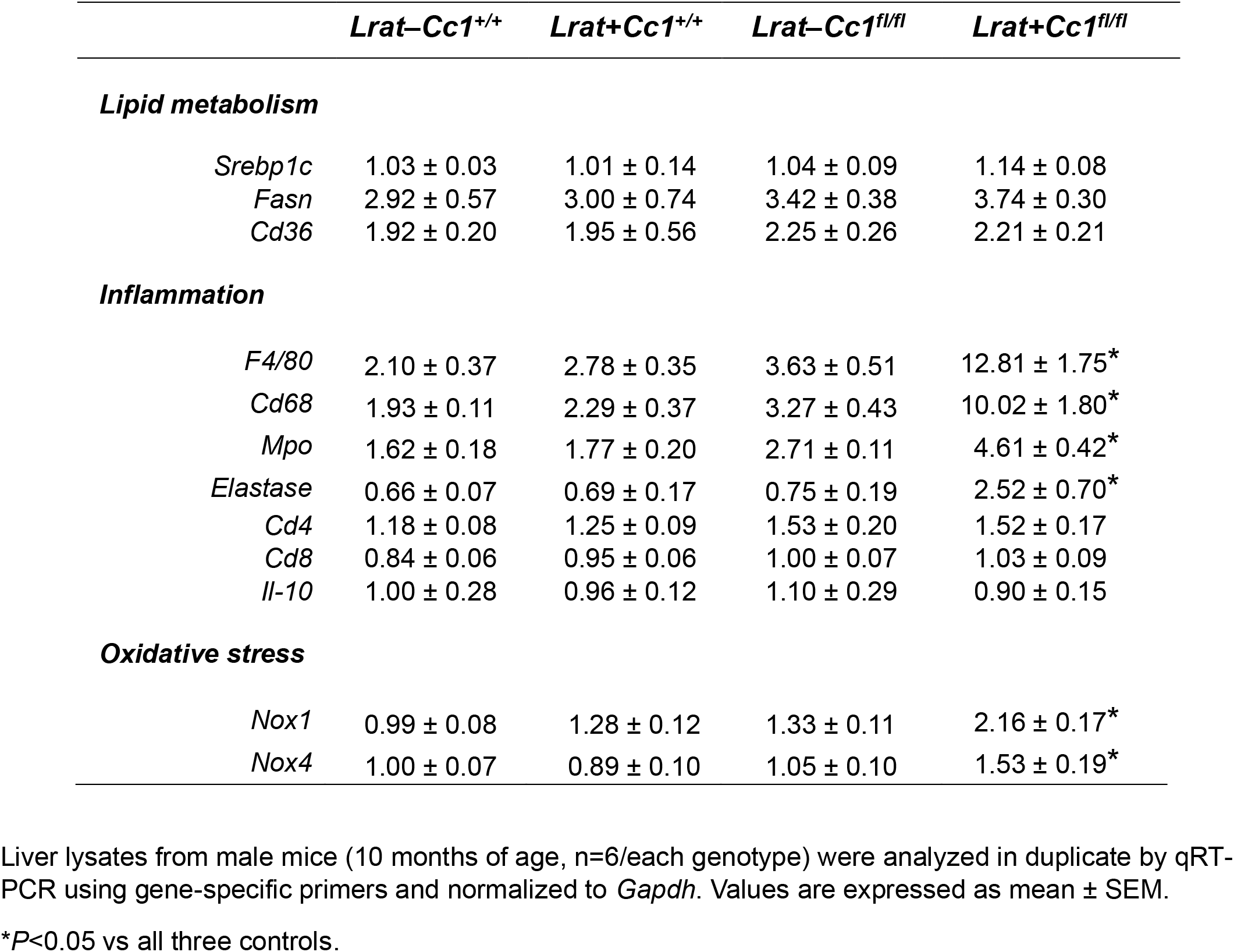
mRNA levels of genes in liver lysates.

**Table S5.**
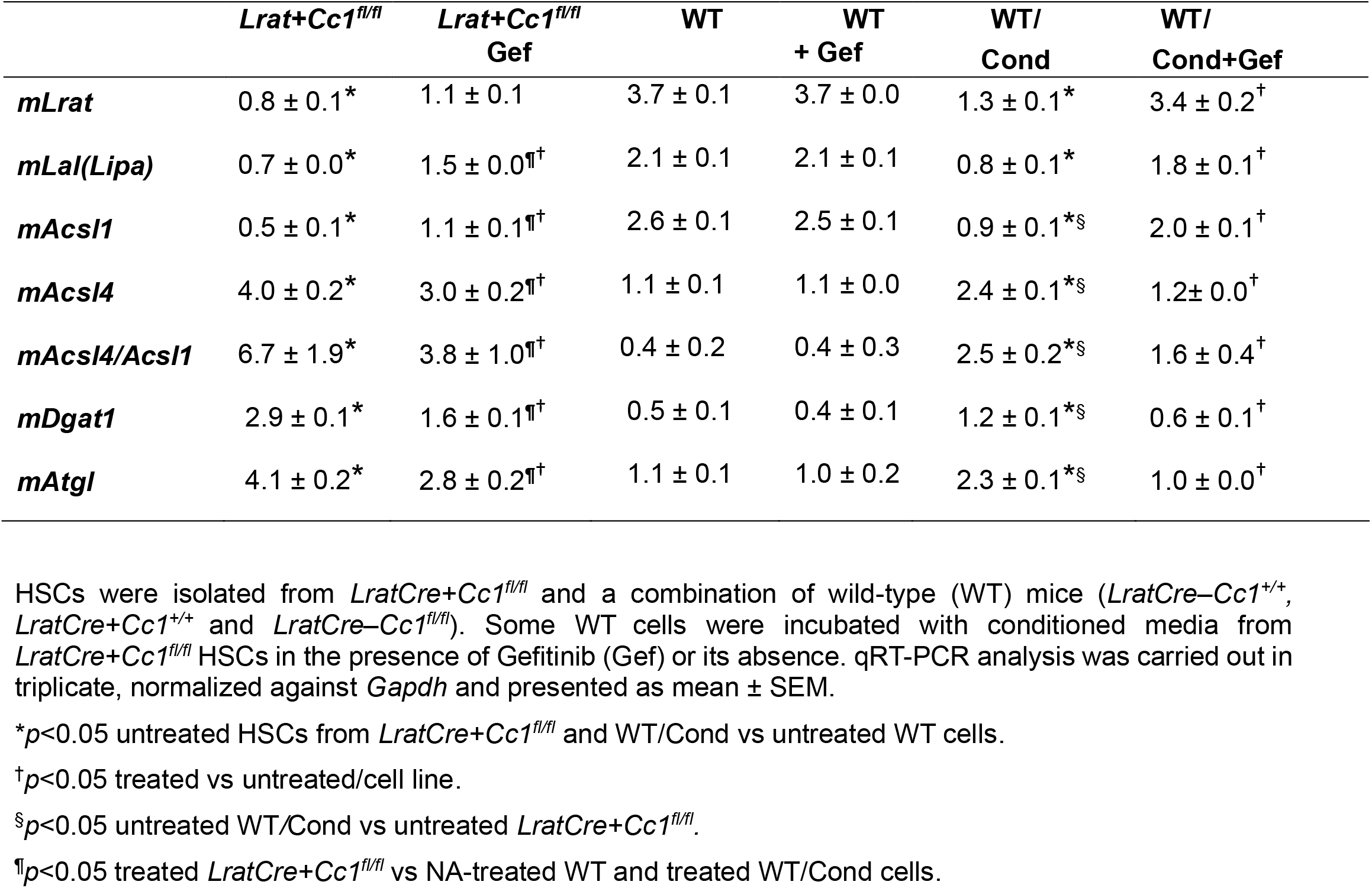
The effect of EGFR inhibitor, Gefitinib, on lipid metabolism of primary HSCs from *LratCre+Cc1^fl/fl^* mice.

**Fig. S1.**
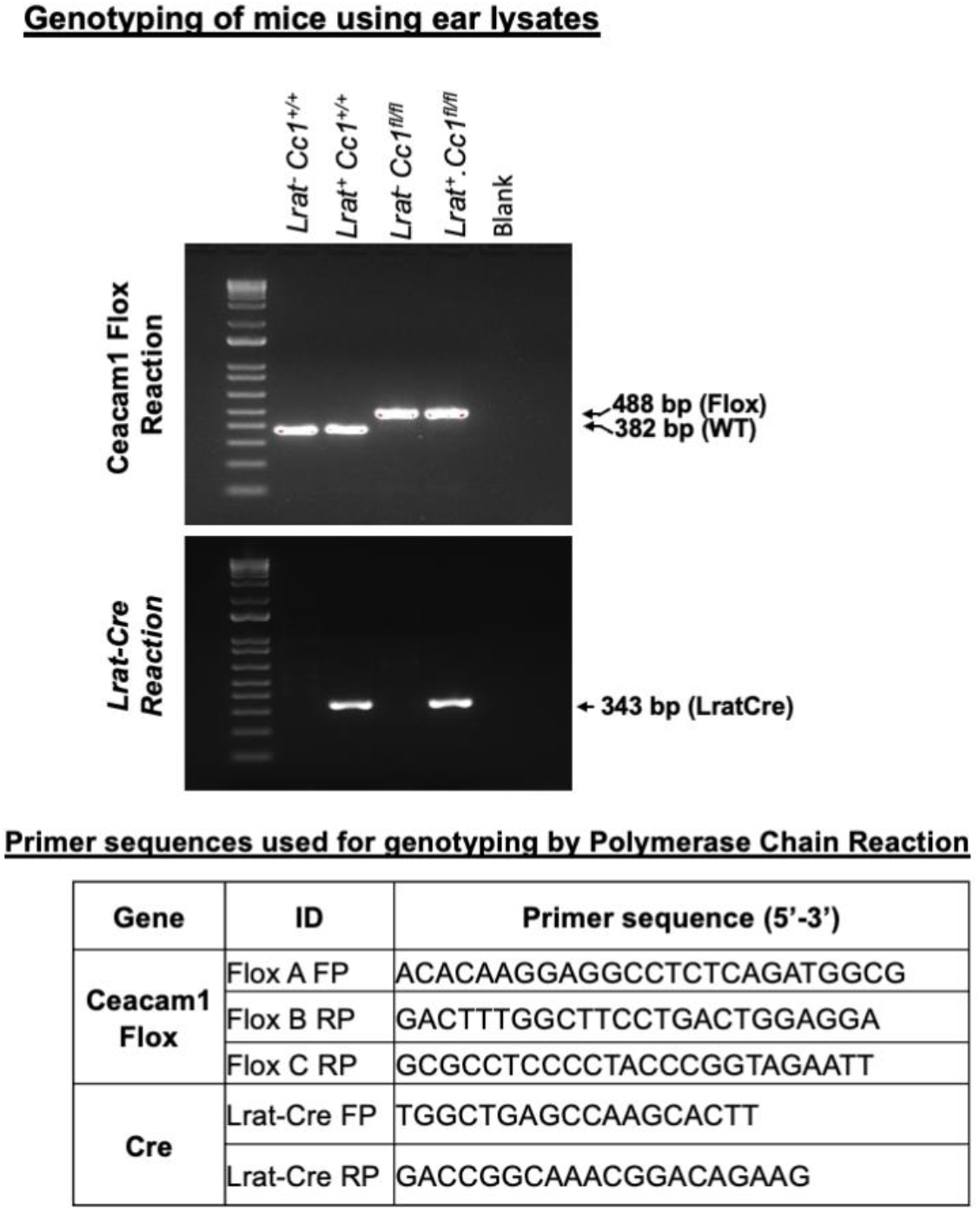
Genotyping of LratCre+Cc1fl/fl mutants. PCR amplification was performed on ear DNA using primers for the FloxA/FloxB pair detected the 382bp wild-type allele (*Cc1^+/+^*) and the FloxA/FloxC pair detected the 488bp null when the primer sets of the Cre reaction were combined. The LratCre+ allele was detected by the 343bp Lrat promoter. Nucleotide sequences were listed in the inserted table. FP denotes forward primer and RP denotes reverse primer.

**Fig. S2.**
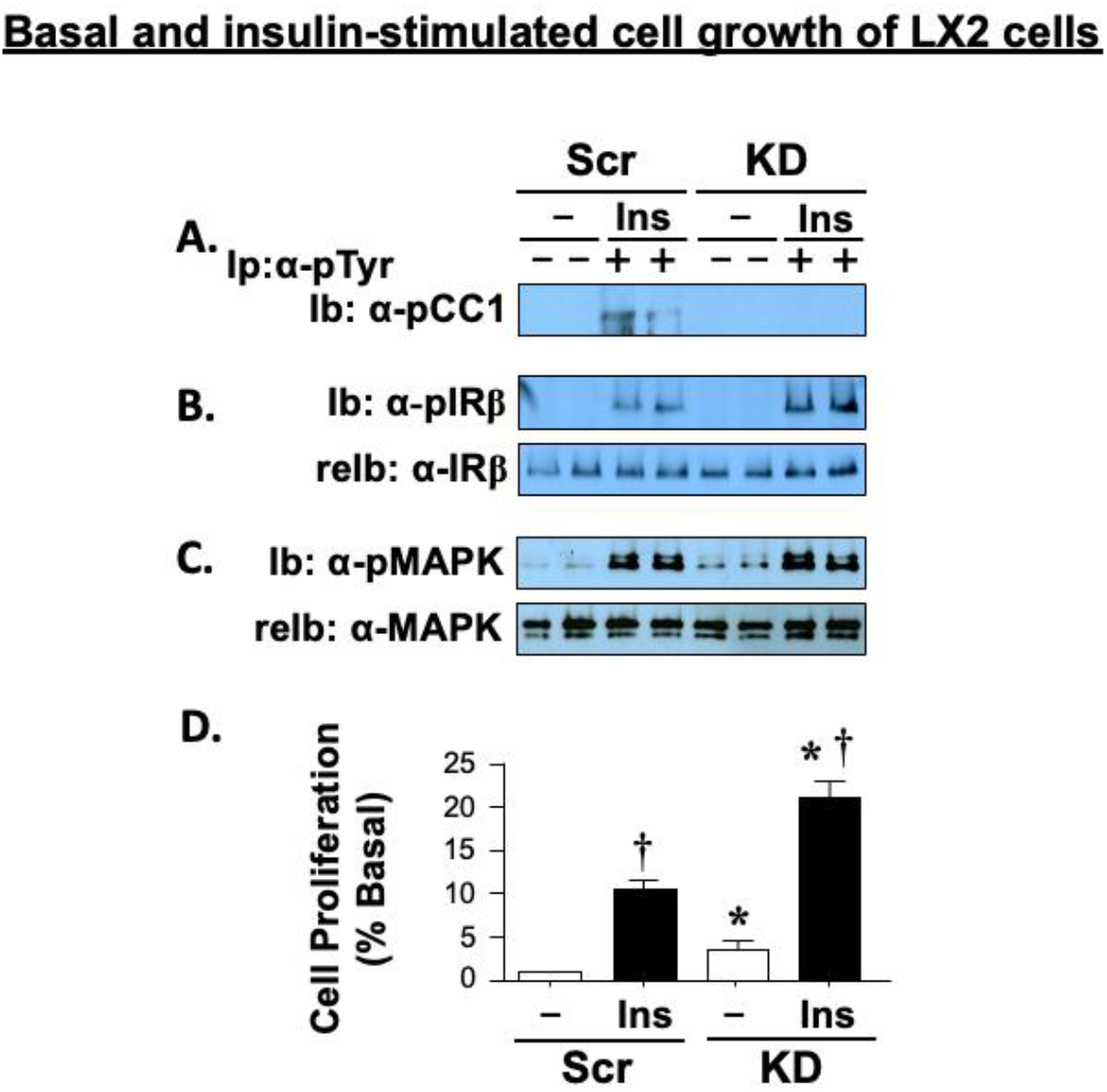
CEACAM1 loss in LX2 cells induces cell growth. Serum-starved Src-LX2 and KD-LX2 cells were treated with insulin (100 nM) for 5 min, lysed and proteins analyzed by immunoblotting (Ib): (A) the α-pTyr immunopellet (Ip) with α-pCEACAM1 (α-pCC1) antibody; (B) total lysates with α-pIRβ followed by re-immunoblotting (relb) with α-pIRβ to normalize for loading; (C) total lysates with α-pMAPK followed by re-immunoblotting (relb) with α-MAPK. (D) Cells were grown in 96 well plates and treated with insulin for 24 hrs before cell growth was assayed in triplicate by MTT assay. Values represent mean ± SEM; *p<0.05 KD-LX2 vs Scr-LX2 cells/treatment group; ^†^p<0.05 insulin-treated vs untreated/cell line.

**Fig. S3.**
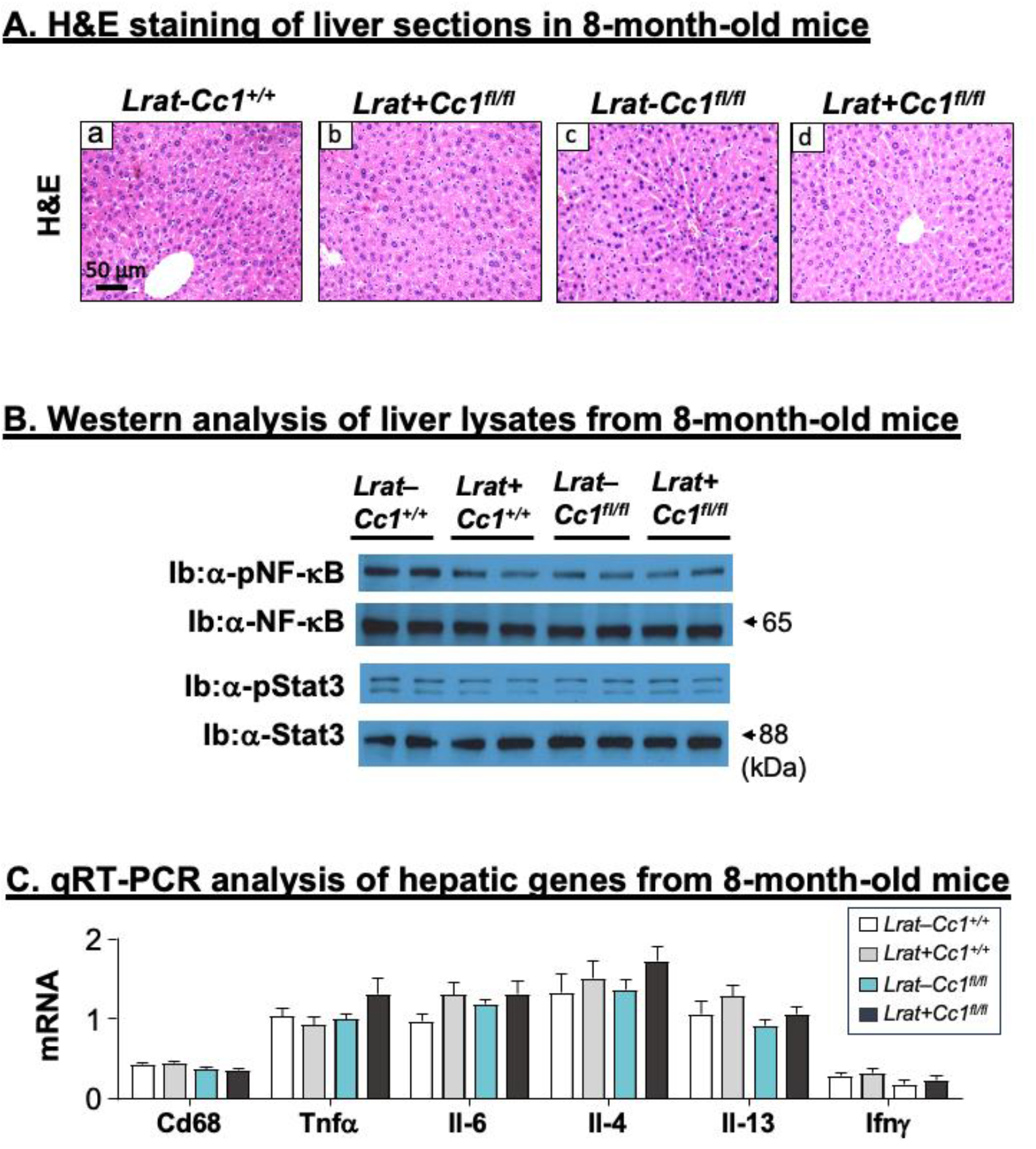
Absence of inflammatory infiltration in 8-month-old mice. Livers were removed from 8-month-old *LratCre+Cc^1fl/fl^* male mice and their 3 littermate controls (n=4–5/genotype) to carry out (A) H&E staining as in the legend of Fig. 5. (B) Western analysis to assess the phosphorylation of the p65 subunit of NF-κB and Stat3 (activation), as in the legend of Fig. 6 and (C) qRT-PCR analysis of the mRNA levels of genes involved in inflammation, as in the legend of Fig. 6.

**Fig. S4.**
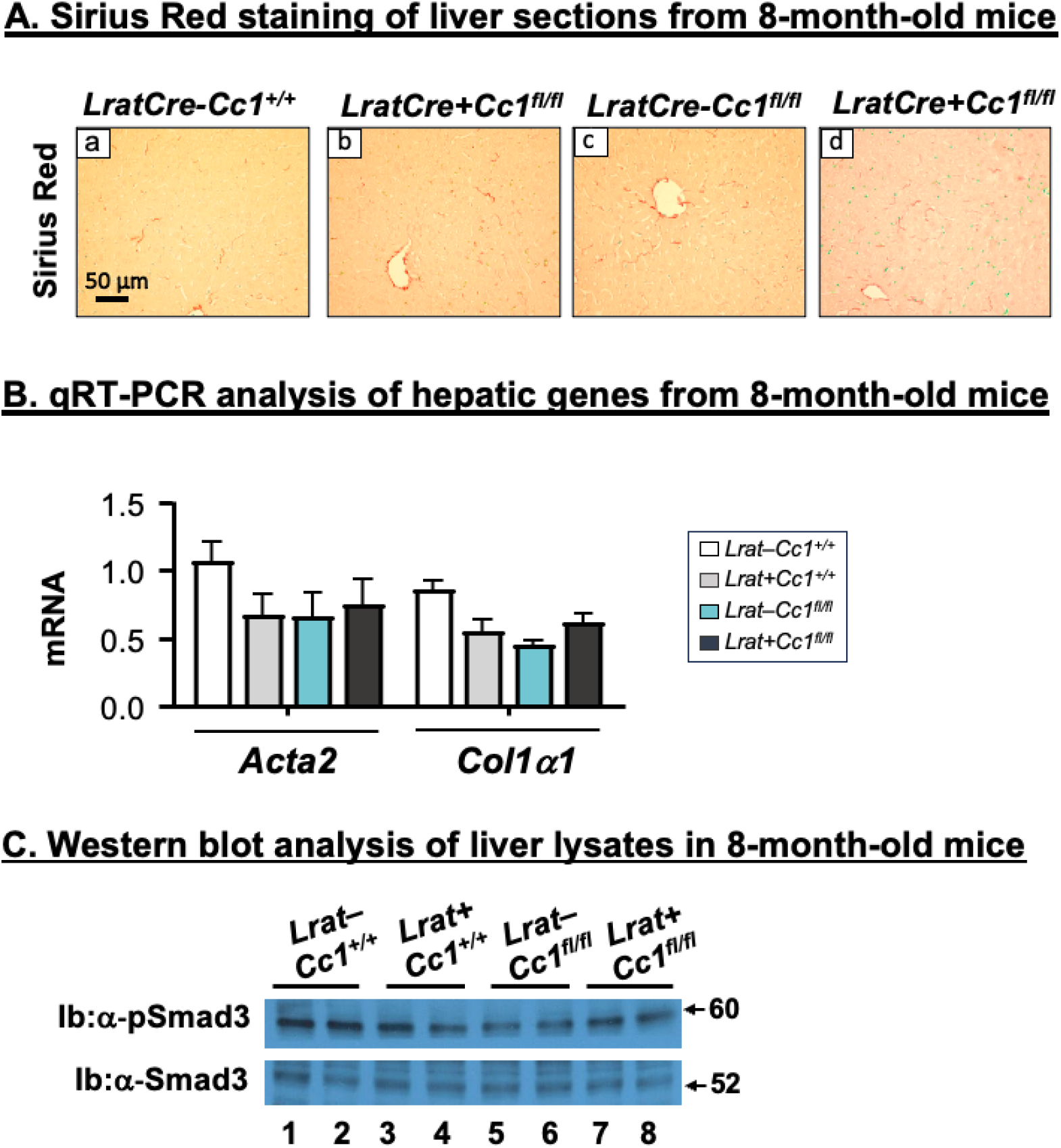
Absence of hepatic fibrosis in 8-month-old mice. Livers were removed from 8-month-old *LratCre+Cc^1fl/fl^* male mice and their 3 littermate controls (n=4–5/genotype) to carry out (A) Sirius red staining of liver sections from mutant mice (panel d) and their littermate controls (panels a-c), as in the legend of Fig. 7. (B) qRT-PCR analysis of the mRNA levels of hepatic genes involved in fibrosis, as in the legend of Fig. 7. (C) Western analysis to assess Smad3 phosphorylation (activation), as in the legend of Fig. 7.

## Notes

### Competing Interest Statement

The authors have declared no competing interest.

